# Nuclear basket proteins Nup2 and Mlp1 drive heat shock-induced 3D genome restructuring downstream of transcriptional activation

**DOI:** 10.1101/2025.01.01.631024

**Authors:** Suman Mohajan, Linda S. Rubio, David S. Gross

**Affiliations:** Department of Biochemistry and Molecular Biology, Louisiana State University Health Sciences Center, Shreveport, LA 71130

**Keywords:** Nuclear Pore Complex, Heat Shock Response, Mlp1, Nup1, Nup2, Nup145, Heat Shock Factor (Hsf1), chromatin, RNA Pol II, transcriptional condensates, 3D genome architecture, *Saccharomyces cerevisiae*

## Abstract

The nuclear pore complex (NPC), a multisubunit complex located within the nuclear envelope, regulates RNA export and the import and export of proteins. Here we address the role of the NPC in driving thermal stress-induced 3D genome repositioning of *Heat Shock Responsive* (*HSR*) genes in budding yeast. We found that two nuclear basket proteins, Nup2 and Mlp1, although dispensable for NPC integrity, are required for driving *HSR* genes into coalesced chromatin clusters, consistent with their strong, heat shock-dependent recruitment to *HSR* gene regulatory and coding regions. *HSR* gene clustering occurs predominantly within the nucleoplasm and is independent of the essential scaffold-associated proteins Nup1 and Nup145. Notably, acute double depletion of Nup2 and Mlp1 has little effect on the formation of Heat Shock Factor 1 (Hsf1)-containing transcriptional condensates, Hsf1 and Pol II recruitment to *HSR* genes, or *HSR* mRNA abundance. Our results define a 3D genome restructuring role for nuclear basket proteins extrinsic to the NPC and downstream of *HSR* gene activation.

## Introduction

The complex three-dimensional (3D) architecture of eukaryotic chromatin is established and maintained through intricate looping and folding interactions. These interactions bring into close proximity enhancers and promoters (that may be tens or hundreds of kilobases apart) as well as promoters and terminators (1–4). In addition to these local looping and folding interactions, chromosomal loci can undergo dynamic repositioning within the nucleus. Such restructuring of 3D chromatin architecture has been suggested to regulate gene expression, replication, DNA repair, chromosomal transposition and mRNA export (5–10). However, the underlying molecular mechanisms by which 3D genome structural changes occur and how these topological alterations impact nuclear processes remain unclear.

Dynamic aspects of 3D chromatin architecture can be investigated using a powerful heat shock-responsive system established in the budding yeast *Saccharomyces cerevisiae*. The heat shock response (HSR) is an evolutionarily conserved, adaptive mechanism that is used by eukaryotic organisms to maintain protein homeostasis in response to thermal, chemical and oxidative stress (11, 12). It is characterized by the transcriptional upregulation of genes encoding heat shock proteins (HSPs) and other homeostasis factors (11, 13) and depends on a sequence-specific transcription factor (TF), Heat Shock Factor 1 (Hsf1), that inducibly binds its cognate heat shock elements (HSEs) situated upstream of *HSR* genes (14–16). We have observed that in response to acute heat shock, Hsf1 forms subnuclear clusters that have characteristics of transcriptional condensates and contain Hsf1, RNA Pol II, and Mediator (17, 18). These condensates drive robust *cis-* and *trans-*intergenic interactions between *HSR* genes (18) that culminate in their coalescence into intranuclear foci (*HSR* gene coalescence [HGC]). While Hsf1, the large subunit of Pol II (Rpb1) and the Mediator Tail subunit Med15 contribute to HGC^1^ (17–19), the potential role of other nuclear factors remains largely unexplored.

It has been previously observed that inducible genes in yeast spatially reposition from the nuclear interior to the periphery upon their activation (20–23). Such repositioning appears to be mediated by the interaction between the genes and the nuclear pore complex (NPC) (24–26), a conserved multiprotein assembly located at the periphery of the nucleus. The yeast NPC is comprised of multiple copies of ∼30 different proteins (nucleoporins (Nups)), 450-500 proteins altogether, which regulate RNA export as well as the import and export of proteins across the nuclear envelope. Nups are either stably or dynamically associated with the NPC (27–30).

The NPC is comprised of outer rings (cytoplasmic ring (CR) and one or two coaxial nuclear rings (NRs)) sandwiched around an inner ring (IR). Each NPC additionally contains a prominent module on its nuclear side termed the nuclear basket (schematically illustrated in Figure 1A). Each outer ring consists of eight Y-shaped heptameric Nup84 complexes (Y-complex) arranged in a head-to-tail orientation around a central axis that passes through the center of the NPC and maintains NPC integrity (29, 31–33). The inner ring is also composed of eight large subunits, commonly referred to as spokes. Together with the coaxial outer rings, the inner ring forms the core scaffold of the complex. This core NPC scaffold connects the nuclear basket, cytoplasmic RNA export complexes, and the membrane rings. The scaffold also anchors the NPC to the nuclear envelope through a small number of transmembrane Nups (29, 32, 34).

**Figure 1.**
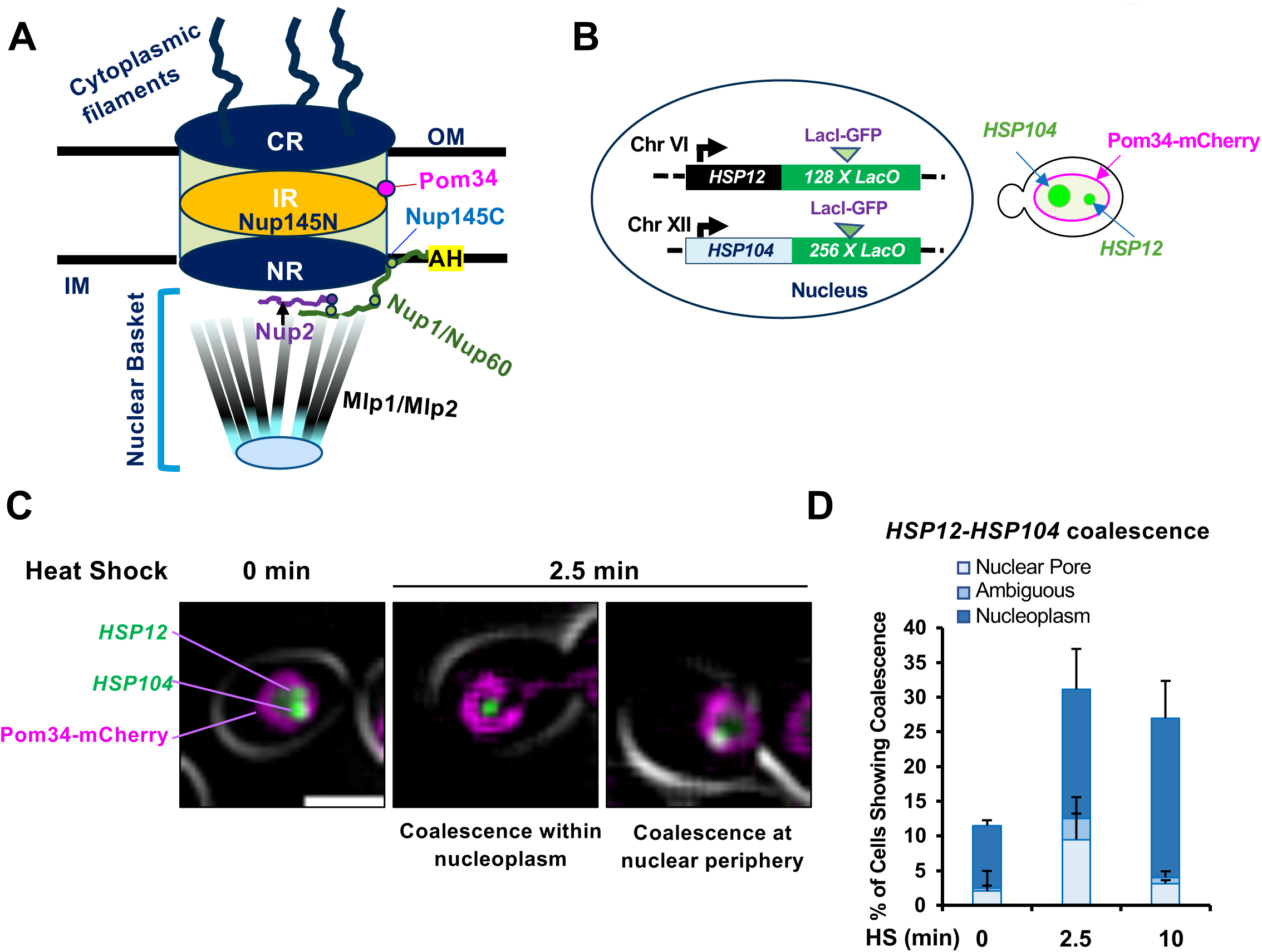
*HSR* gene coalescence occurs both within the nuclear interior and at the nuclear periphery. A. The yeast NPC. IM, Inner membrane; OM, Outer membrane; CR, Cytoplasmic Ring; IR, Inner Ring; NR, Nuclear Ring; AH, Amphipathic helix (present in Nup1, through which it anchors to the nuclear membrane); Nup145C, C-terminal domain of Nup145 (part of the Y complex, a CR and NR component); Nup145N, N-terminal domain of Nup145 (located within the central tube of the NPC). Each Nup exists in multiple copies; only one representative copy is shown. B. Schematic depiction of the LacI-GFP tagged heterozygous diploid strain ASK706 bearing one copy each of *HSP104-lacO_256_* and *HSP12-lacO_128_* and expressing the mCherry-marked nuclear pore transmembrane protein Pom34. C. Coalesced *HSP104* and *HSP12* gene foci are found at the NPC and within the nucleoplasm. ASK706 cells were imaged both prior to and following a 25° to 38°C heat shock (HS) on a widefield fluorescence microscope. Two examples of 2.5 min HS, with different localization for the coalesced foci, are depicted. Scale bar: 2 µm. D. Quantification of *HSP104-HSP12* localization data, scored using the nuclear pore marker Pom34-mCherry, in cells treated as in C. On average, 100 cells were scored per sample. N=2. Error bars represent standard deviation.

Nup145 is an essential nucleoporin that undergoes autoproteolytic cleavage and generates two functionally distinct polypeptides, Nup145C and Nup145N (orthologs of human NUP96 and NUP98, respectively). Nup145C integrates into the Y complex, acts as a scaffold and is slowly exchanging, while Nup145N becomes a part of the central core, is more dynamic and can link different NPC modules (35, 36). In addition, Nup145 has been suggested to associate with chromatin and assist in the repositioning of an activated gene to the NPC (37).

The nuclear basket consists of FG or FXF(G) domain-containing Nups (Nup1, Nup2, and Nup60) and coiled-coil domain-containing Nups (Mlp1 and Mlp2) (33, 36, 38–41). While Nup1 and Nup60 are scaffold-associated and connect the basket to the nuclear membrane (33, 42), the filamentous Mlps extend toward the nucleoplasm and converge into a distal ring (29, 33, 38, 40, 43) (see Figure 1A). It has been suggested that the nuclear basket regulates membrane curvature and NPC integrity, distribution, and mobility (30, 40, 42). Additionally, the nuclear basket serves as a docking site for mRNP granules, facilitating mRNA quality control and efficient export (44). In addition to its canonical transport function, the nuclear basket is also involved in DNA repair, transcriptional regulation and gene targeting (44–49). How the nuclear basket regulates these processes and chromatin structures remains poorly understood. Key roles of Mlp1, Nup1, Nup2 and Nup145 are summarized in Table 1.

**Table 1.** Role of NPC proteins.

| Features | Nup2 | Mlp1 | Nup1 | Nup145 | References |
| --- | --- | --- | --- | --- | --- |
| FG repeats | Yes | No | Yes | Yes (Nup145N)<br>No (Nup145C) | (102–104) |
| Phase separation propensity | Yes (PSP score 0.9453) | Yes (PSP score 0.8533) | Yes (PSP score 0.9772) | Yes (PSP score 0.7217) | PSPredictor(105) |
| Nonessential/ Essential | Non-essential | Non-essential | Essential | Essential | SGD (yeastgenome.org) |
| mRNA export | Yes /Mild | Yes /Mild | Yes | Yes | (63, 67, 68, 86, 87, 106–110) |
| mRNA quality control | ? | Yes | ? | ? | (67, 88) |
| NPC and NE integrity | ? | ? | Yes | Yes | (42, 60, 63) |
| NPC components | NPC Basket | NPC Basket filament | Connector of NPC basket to NPC and NE | Nup145C- Scaffold of NPC, Nup145N- NPC central core | (33, 34, 36, 38–42) |
| Dynamic /stable | Dynamic | Dynamic | Dynamic | Nup145N- Dynamic<br>Nup145- Very slow exchange | (36, 39, 40, 111). |
| Insulator/ boundary activity | Yes | ? | ? | ? | (89, 90) |
| Role in repositioning of inducible genes to NPC | Yes (SSA2, SSA4, HIS4; GAL1-10; INO1, HAS1- TDAI | Yes (α-factor-induced genes, HXK1; GAL1-10, HSP104) | Yes (INO1, GAL10) | Yes SUC2 | (5, 22, 37, 46, 47, 57, 61, 62, 66, 70, 91, 110, 112–114) |
| Chromatin association (ChIP, IF, CHEC) | Yes (GAL genes, HXK1, INO1) | Yes (GAL2, GAL1-10), HXK1, INO1) | ? | Yes (SUC2) (Nup145C) | (5, 22, 24, 37, 46, 47, 70, 71, 91, 112) |

The NPC has been implicated in regulating transcription in yeast (20, 21, 46). Likewise, NPC proteins have been found associated with mammalian enhancers and super-enhancers (SEs) (8, 50–53) and Mlp1/2 and Nup145N orthologues in *Drosophila* are particularly enriched in active chromatin and may foster formation of enhancer-promoter loops (54, 55). It has been suggested that certain nucleoporins can undergo phase separation that may facilitate the formation of transcriptional hubs (53). Such hubs then concentrate chromatin structural proteins and transcriptional coactivators at the SEs that may facilitate transcription or mRNA export or both. Coalescence of *HSR* genes in yeast bears important similarities to mammalian super-enhancers, including the presence of extensive DNA loops and transcriptional condensates that concentrate chromatin-associated TFs and transcriptional machinery and co-activators (18, 56). Here we provide evidence that in response to acute heat stress, *HSR* genes coalesce at the NPC as well as within the nucleoplasm. While the essential scaffold protein Nup145 and scaffold-associated nuclear basket protein Nup1 play no detectable role in *HSR* gene coalescence, the dynamically exchangeable nuclear basket proteins Mlp1 and Nup2 play a critical role and likely do so in their NPC-free state.

## Results

### Coalescence of *Heat Shock Response* genes preferentially occurs within the nucleoplasm

Previous studies have suggested that a variety of inducible genes in *S. cerevisiae,* including those that respond to stress, relocate from the nuclear interior to the nuclear periphery upon their transcriptional activation as discussed above. Certain of these genes have a Gene Recruitment Sequence (GRS) within their upstream regions that is implicated in such repositioning (24–26, 57). However, the underlying molecular mechanisms remain largely unknown. We have previously reported that Hsf1-regulated *HSR* genes, dispersed across multiple chromosomes, physically interact and cluster together within minutes following cell exposure to proteotoxic stresses such as heat shock or ethanol (17–19, 58).

To address whether such coalescence occurs at the nuclear pore or within the nucleoplasm, we constructed a strain in which the nuclear pore transmembrane protein Pom34 was labeled with mCherry and the *HSP104* and *HSP12* genes were tagged with *LacO* arrays to which LacI-GFP binds (schematically summarized in Figure 1B). We then conducted live imaging of cells subjected to an instantaneous HS (25°C to 38°C) and examined coalescence of the two genes following 2.5 and 10 min of HS. As shown in Figure 1C, coalescence occurred at both the nuclear pore and within the nucleoplasm. Overall, of the cells that exhibited *HSP12-HSP104* coalescence (∼30% of the total), coalescence at the periphery was observed in less than one-third, while coalescence within the nucleoplasm was observed in greater than two-thirds (Figure 1D). Fixed cell microscopy similarly revealed examples of HS-induced *HSP12-HSP104* coalescence at the nuclear periphery as well as within the nuclear interior^2^. These data indicate that *HSR* gene coalescence can occur within the nucleoplasm, and it is likely the preferred location.

### Nup1 and Nup145 are important for maintaining integrity of the NPC but play no detectable role in heat shock-induced *HSR* gene coalescence

As discussed above, certain NPC proteins have been implicated in the repositioning of inducible genes to the nuclear pore upon their activation. To address whether the NPC is required to drive the coalescence of *HSR* genes, we initially tested the involvement of two essential, scaffold-associated nucleoporins, Nup1 and Nup145. To do so, we conditionally depleted these proteins using the Auxin Inducible Degradation (AID) technique (59). Both proteins have been reported to maintain integrity of the NPC and nuclear envelope (42, 45, 60). They also have been implicated in repositioning activated genes to the nuclear pore (37, 57, 61, 62). We tagged each gene with a mini-degron (Figure 2A) and as shown in Figure 2B, 85 - 95% degradation of each protein was achieved within 60 min of adding auxin.

**Figure 2.**
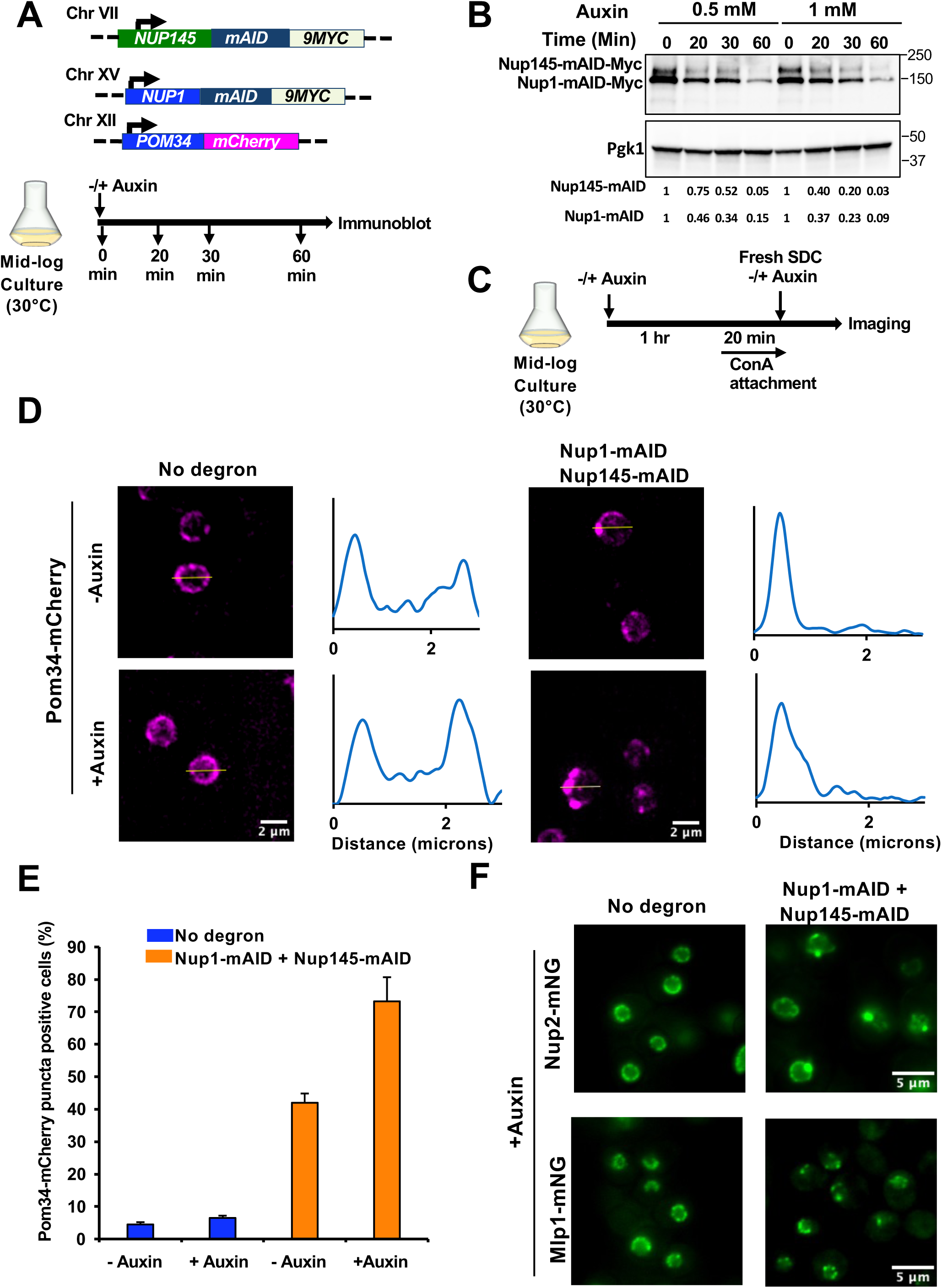
Simultaneous depletion of Nup1 and Nup145 perturbs NPC integrity and alters the distribution of Nup2 and Mlp1. (A) Top: Haploid strain SMY163 bears *NUP145-mAID-9MYC*, *NUP1-mAID-9MYC* and *POM34-mCherry.* Bottom: Experimental strategy for optimizing auxin concentration and incubation time for double degradation. (B) Immunoblot analysis of Nup1-mAID-9Myc and Nup145-mAID-9Myc in cells treated with different concentrations of indoleacetic acid (IAA) for the indicated times. Monoclonal antibody 9E10 was used to detect the levels of the respective proteins. Endogenous Pgk1 was used as a loading control. (C) Experimental strategy for assessing NPC integrity upon simultaneous depletion of Nup1 and Nup145. Cells expressing mCherry-tagged Pom34 that bear either no degron (SMY160) or the Nup1-, Nup145-double degron (SMY163) were pretreated with 1 mM IAA or vehicle alone (0.17% ethanol) for 60 minutes at 30°C, followed by attachment to a ConA-coated VAHEAT substrate. Imaging was done using an Olympus W1 fluorescence spinning disk confocal microscope (see Methods). (D) Subnuclear localization of Pom34-mCherry in cells treated (or not) with 1 mM IAA. A representative maximum projection image (21 z-planes with a step size of 0.3 microns) of each treatment is shown. The intensity profile plot for each representative image is displayed on the Right. (E) Quantification of cells showing NPC punctate structure at the nuclear periphery in the indicated strains. Depicted values are means + SD, N=2. 100 cells per biological sample were analyzed. (F) Subcellular localization of Nup2-mNeonGreen (mNG) and Mlp1-mNG in no-degron strains (SMY216 and SMY192) and double-degron strains (SMY221 and SMY196) maintained at 25°C and treated with 1 mM IAA for ∼60 min prior to imaging on a widefield fluorescence microscope. A representative single z-plane is shown for each sample (step size of 0.5 microns).

To examine the effect of simultaneous depletion of Nup1 and Nup145 on NPC integrity, we performed imaging of live cells expressing mCherry-labeled Pom34 (Figures 2C, 2D), a protein associated with the NPC inner ring (29). NPC integrity was compromised upon double depletion of Nup1 and Nup145, as ∼75% of cells harbored at least one Pom34 punctate structure (Figure 2D, 2E). Note that auxin alone did not affect NPC integrity, while the degron tag itself had some impact. This observation is consistent with previous studies that found a mutation in either the N- or C-terminal domains of Nup145 compromised NPC structure (31, 60, 63). We additionally tested the effect of simultaneously depleting Nup1 and Nup145 on the integrity of the nuclear basket. Consistent with its effect on Pom34, double deletion of Nup1 and Nup145 disrupted the subnuclear localization of both Nup2-mNeonGreen (mNG) and Mlp1-mNG, leading to the formation of punctate structures (Figure 2F).

To investigate the effect of Nup1-, Nup145-double depletion on the physical clustering of *HSR* genes (*HSR* gene coalescence), we used a powerful crosslinking, digestion and proximity ligation-based technique, Taq I - 3C (19, 64). Cells were treated with auxin and then subjected to an acute HS as above. Previous work has shown that *HSR* gene clustering peaks rapidly (within 2.5 - 10 min) (58); here we conducted 3 min HS. 3C analysis of the parental strain lacking a degron-tagged gene revealed a dramatic HS-dependent increase in both *cis-* and *trans-* intergenic interactions (Figures 3A and 3B), as previously observed (17–19, 58, 65). (See Figure S1 for *HSR* gene physical maps.) Also as previously observed, there was almost no detectable 3C signal in non-stressed (0 min HS) cells. Notably, double depletion of Nup1 and Nup145 had little or no impact on the HS-induced interactions (Figure 3A, 3B; compare orange vs. blue bars). Collectively, these results indicate that while Nup1 and Nup145 are important for maintaining NPC integrity and the wild-type distribution of nuclear basket proteins, they play little or no role in driving the 3D restructuring of *HSR* genes following heat shock. Moreover, this observation is congruent with the idea that *HSR* gene coalescence predominantly occurs extrinsic to the NPC, as suggested above.

**Figure 3.**
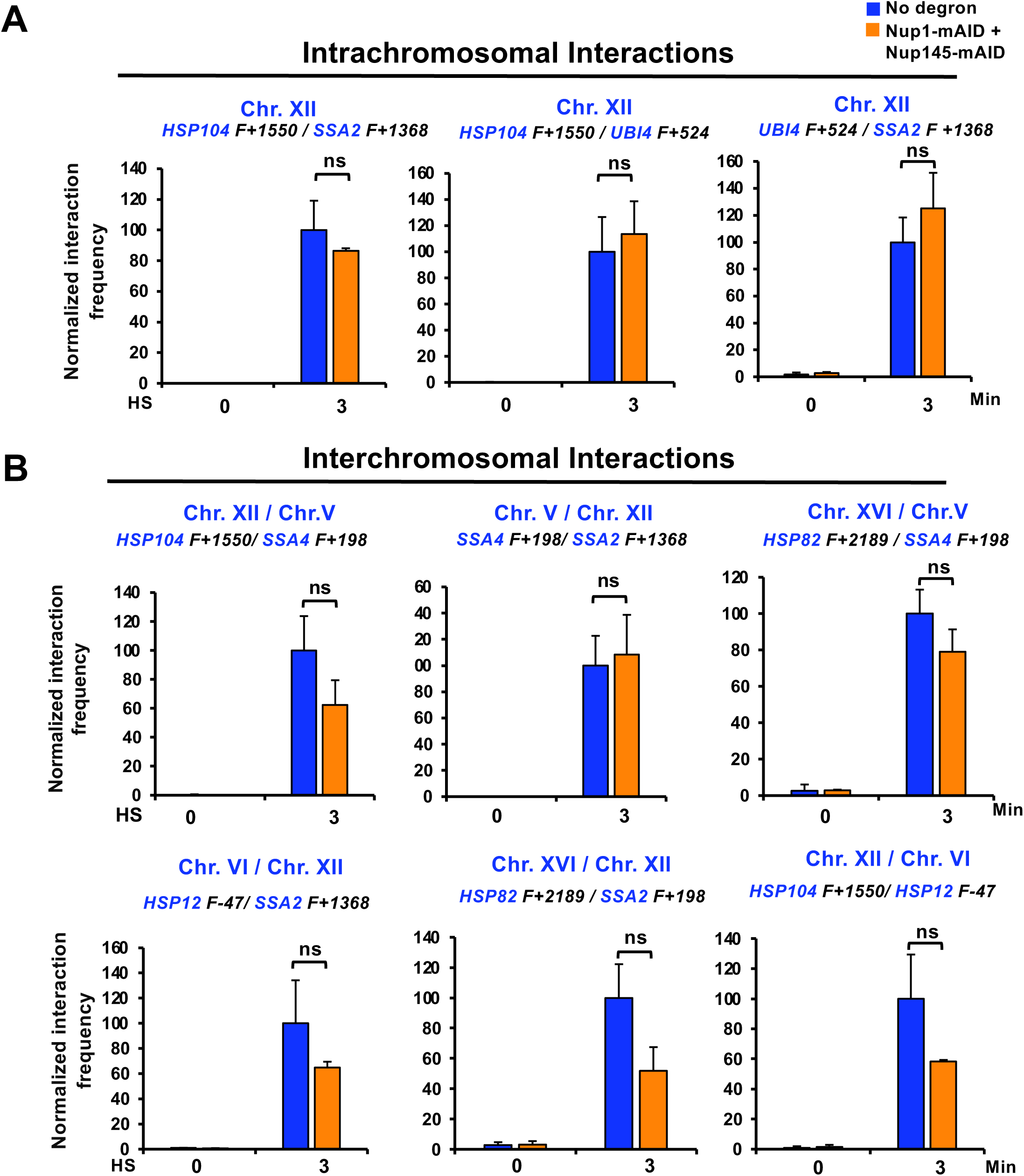
Nup1-, Nup145-double depletion has minimal effect on heat shock-induced *HSR* intergenic interactions. (A) Frequency of *cis-*interactions between the indicated Hsf1-regulated genes as detected by TaqI-3C. No degron (LRY016) and double degron (SMY148)-tagged cells were pretreated with 1 mM IAA for 60 min at 30°C prior to exposing them (or not) to HS (39°C). Primer locations for loci examined in this study are illustrated in Figure S1. Pairwise tests used forward (F; sense strand identical) primers positioned nearby the indicated Taq I site. Interaction frequencies were normalized with respect to the control strain, which was arbitrarily assigned 100. Shown are means + SD, N=2, qPCR=4. To determine significance, an unpaired two-tailed t-test was performed. ns, not significant (*p>0.05*). (B) As in A, except *trans*-interactions were assayed.

### Nup2 and Mlp1 are critically required to drive *HSR* genes into coalesced chromatin clusters

Next, given their reported role in targeting transcriptionally active genes to the NPC (46, 47, 66), we wished to investigate what role, if any, the nuclear basket proteins Nup2 and Mlp1 have in driving *HSR* gene coalescence. To address this, we performed live cell imaging of fluorophore-tagged genes and quantitative 3C of untagged and degron-tagged strains. For live cell imaging, we used heterozygous diploids that harbored single alleles of LacI-GFP marked *HSP104-lacO_256_* on Chr. XII and tetR-mCherry marked *HSP82-tetO_200_* on Chr. XVI (Figure 4A) and bore homozygous deletions of *MLP1*, *NUP2* or both. While chronic exposure to elevated temperature caused a fitness defect in both *nup2Δ / nup2Δ* and *mlp1Δ / mlp1Δ* cells, brief exposure had no effect on cell viability (Figures S2A and S2B). Cells were subjected to instantaneous HS (38°C) for various durations (0, 2.5, 10, and 17.5 min), and images were captured using widefield fluorescence microscopy. As shown in Figure 4B, deletion of *NUP2* or *MLP1* alone had only a mild effect on *HSP104* - *HSP82* gene coalescence. In marked contrast, double deletion of *NUP2* and *MLP1* significantly reduced the frequency of *HSP104-HSP82* interaction (Figure 4C).

**Figure 4.**
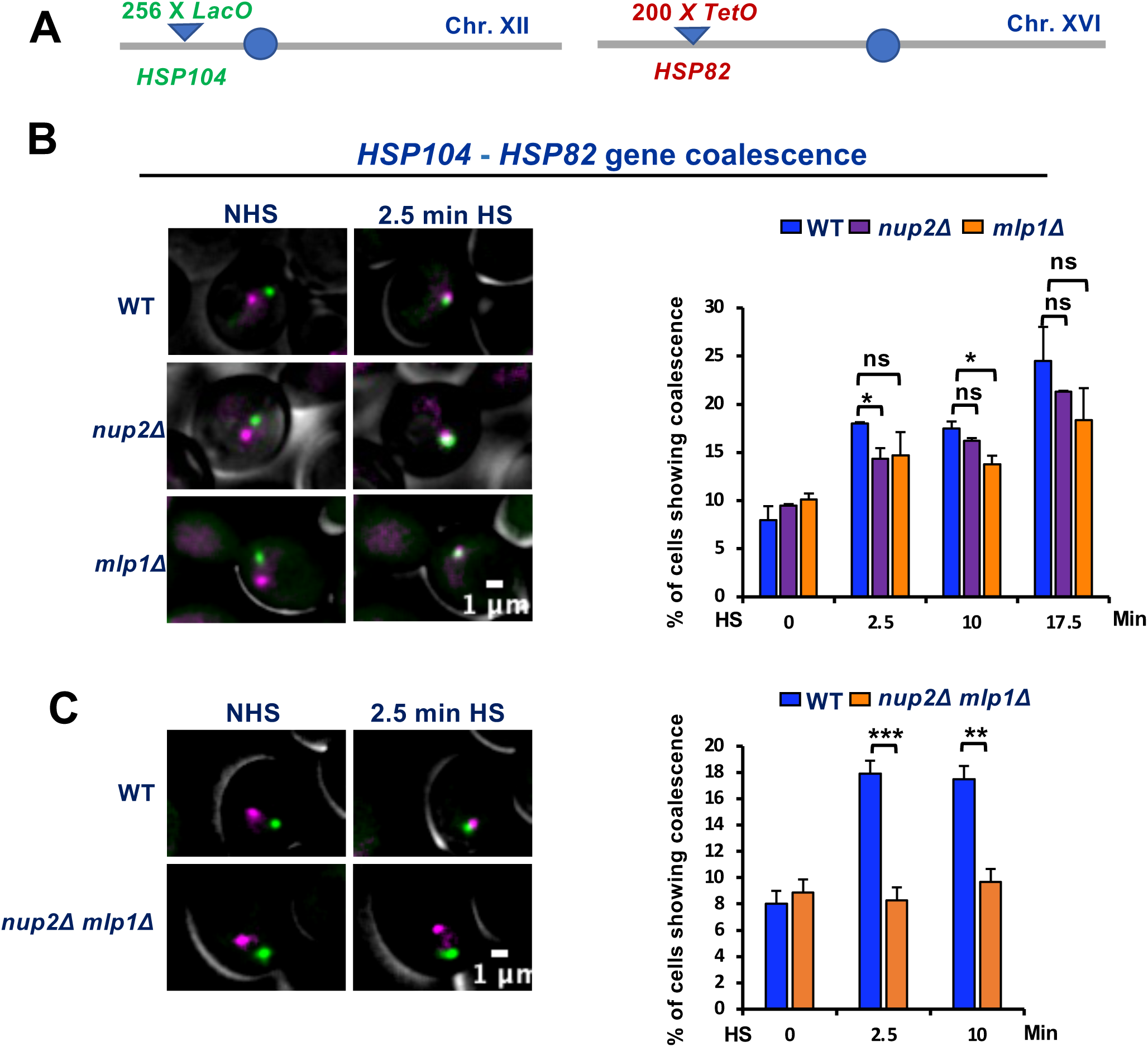
Deletion of either *NUP2* or *MLP1* has only a mild effect on heat shock-induced *HSR* gene interactions while deletion of both significantly reduces the frequency of *HSR* gene coalescence. (A) Chromosomal location of *HSP104-LacO* and *HSP82-TetO* alleles in SMY206 and its derivative diploid strains. All strains ectopically express LacI-GFP and tetR-mCherry fusion proteins. (B) Live cell imaging of SMY206 (WT), SMY208 *(nup2Δ/nup2Δ)* and SMY207 (*mlp1Δ/mlp1Δ)* cells both prior to and following an instantaneous 25°C to 38°C heat shock. Illustrated are representative images within the Z stack showing the intracellular location of *HSP104* (LacI-GFP labeled) and *HSP82* (tetR-mCherry labeled). *Right:* Quantification of cells exhibiting *HSP104*-*HSP82* gene coalescence in the indicated strains and at the indicated times. Approximately 100 cells were scored per biological sample per time point. Statistical significance was determined using an unpaired two-tailed t-test. \**p*<0.05; ns, not significant. Depicted values are means + SD, N=2. (C) Live cell imaging of SMY206 (WT) and LRY119 *(nup2Δ/nup2Δ mlp1Δ/mlp1Δ)* cells treated, illustrated and quantified as above. An average of 60 cells was analyzed per biological replicate per time point. Statistical significance was assessed as above.

As an orthogonal approach, we constructed Nup2-mAID and Mlp1-mAID expressing strains (Figure 5A) and performed 3C. Following addition of auxin for 30 min, 90 to 95% degradation of degron-tagged Nup2 and Mlp1 was achieved (Figures 5B, 5C). As shown in Figure S3A, depletion of either Nup2 or Mlp1 had a mild effect on both long-range *cis* (intrachromosomal) and *trans* (interchromosomal) *HSR* intergenic interactions, consistent with the above microscopic analysis of single homozygous mutants. We additionally tested the role of Nup2 and Mlp1 on HS-induced intragenic interactions. In previous work, we observed that transcriptionally activated *HSR* genes not only engage in intergenic interactions but also engage in intragenic looping between enhancer-promoter (E-P), promoter - coding region, and promoter - 3’-UTR (17–19, 58). Depletion of either Nup2 or Mlp1 alone had, for the most part, only a modest effect on such interactions (Figure S3B). These results suggest that Nup2 and Mlp1 individually have a limited non-redundant role in driving either the spatial repositioning or intragenic restructuring of *HSR* genes in response to heat shock.

**Figure 5.**
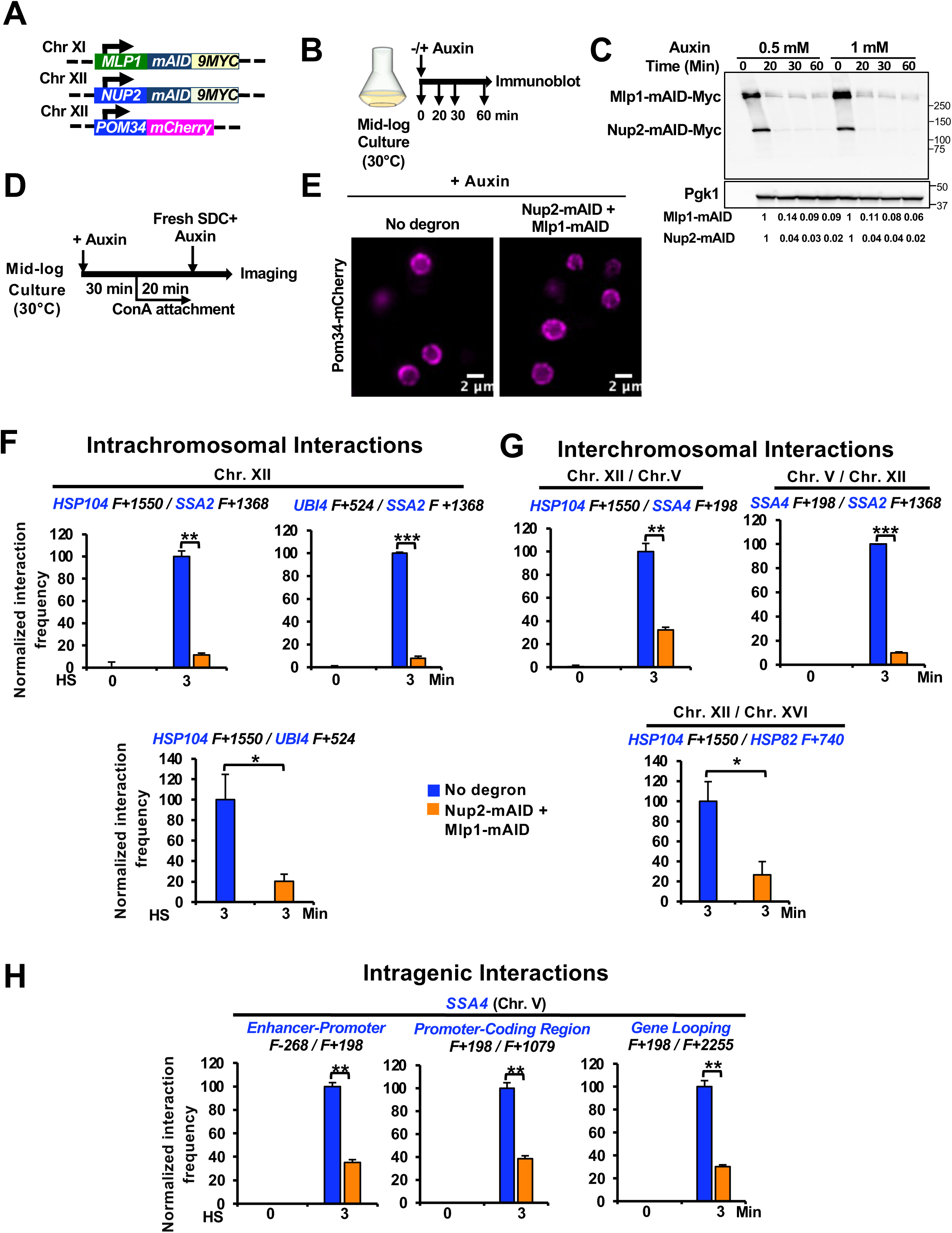
Double depletion of Nup2 and Mlp1 has no detectable effect on NPC integrity but significantly reduces heat shock-induced *HSR* intergenic and intragenic interactions. (A) Relevant genotype of strain SMY182, a haploid strain bearing *MLP1-mAID-9MYC, NUP2-mAID-9MYC* and *POM34-mCherry*. (B) Experimental strategy for optimizing auxin concentration and incubation time. (C) Immunoblot analysis of Mlp1-mAID-9Myc and Nup2-mAID-9Myc in cells treated with different concentrations of IAA for the indicated times. Detection and load control were as in Figure 2B. (D) Experimental strategy for assessing NPC integrity upon simultaneous depletion of Mlp1 and Nup2. Cells expressing mCherry-tagged Pom34 bearing either no degron (SMY160) or the Mlp1-, Nup2-double degron (SMY182) were pretreated with 0.5 mM IAA at 30°C for 30 minutes, followed by attachment of cells on ConA-coated coverslips and imaging using widefield fluorescence microscopy (see Methods). (E) Subnuclear localization of mCherry-tagged Pom34. Images were captured across 11 z planes with a step size of 0.5 microns using widefield fluorescence microscopy. A representative plane of the Z stack is shown for each condition. (F) & (G) Taq I-3C assay depicting *cis-* and *trans-*interactions between Hsf1-regulated genes in no degron (LRY016) and double degron (SMY152) strains. Cells were pretreated at 30°C with 0.5 mM of IAA for 30 min and then incubated at 25°C for 10 min (NHS) or upshifted from 25°C to 39°C for 3 min (HS) prior to crosslinking. All other steps and data presentation are as in Figure 3. Shown are means + SD, N=2, qPCR=4. To determine significance, an unpaired two-tailed t-test was performed. *\*, p<0.05; **, p<0.01, ***, p<0.001*. (G) Intragenic interactions detected within the *SSA4* locus analyzed as above.

Given that the nuclear basket regulates the distribution and mobility of the NPC (30) as well as NPC and nuclear envelope integrity (42), we wondered if together, Nup2 and Mlp1 might play a more substantial role in maintaining either NPC integrity or in governing 3D genome topology in heat-shocked cells. To address this, we performed a conditional double depletion of Nup2 and Mlp1 using the AID strategy as above and initially investigated its effect on NPC integrity by examining subnuclear localization of mCherry-tagged Pom34 (Figures 5A, 5D). Live cell imaging revealed that Pom34 perinuclear localization remained unaffected (Figure 5E), consistent with the notion that Nup2 and Mlp1 are not critical for NPC integrity.

To address the combined impact of Nup2 and Mlp1 on 3D genome architecture, we performed quantitative 3C on HS-induced no-degron and double-degron cells that had been previously exposed to auxin as above. As shown in Figures 5F and 5G, simultaneous depletion of Nup2 and Mlp1 substantially reduced both intrachromosomal and interchromosomal interactions between *HSR* genes. These results are consistent with the *HSP104-HSP82* coalescence phenotype of the *nup2Δ mlp1Δ* double deletion strain (Figure 4C). In both cases, removal of both Nup2 and Mlp1 strongly suppressed the spatial repositioning of Hsf1 target genes. The combined depletion of Nup2 and Mlp1 not only impaired long-range intergenic interactions but also short-range intra-locus interactions (Figures 5H and S4). Notably, double depletion of Nup2 and Mlp1 had a minimal effect on the viability of acutely heat-shocked cells (Figure S2C). Altogether, these results indicate that the dynamic nuclear basket proteins are dispensable for maintaining NPC integrity, yet they have important, partially redundant roles in driving the heat shock-dependent 3D reorganization of *HSR* genes. In light of this striking observation, we focused on Nup2 and Mlp1 in the remaining analyses.

### In thermally stressed cells, Nup2 and Mlp1 relocalize within the nucleus and are inducibly recruited to *HSR* genes

To gain insight into how Mlp1 and Nup2 impact 3D genome structure, we investigated the intranuclear location of each protein following exposure to acute HS. We tagged them with mNG and as shown in Figures 6A and 6B, both proteins, enriched at the nuclear periphery in the non-heat shock (NHS) state, rapidly relocalized in cells exposed to HS, and this was evident as early as 2.5 min. Under the control condition, Mlp1 showed a discontinuous pattern, consistent with the fact that NPCs positioned near the nucleolus typically lack Mlp1/2 (67, 68). Following HS, both Nup2 and Mlp1 became diffusely distributed throughout the nucleoplasm, with a fraction of Mlp1 molecules coalescing into several discrete intranuclear foci (Figure 6A). The latter observation is consistent with previous reports (68, 69).

**Figure 6.**
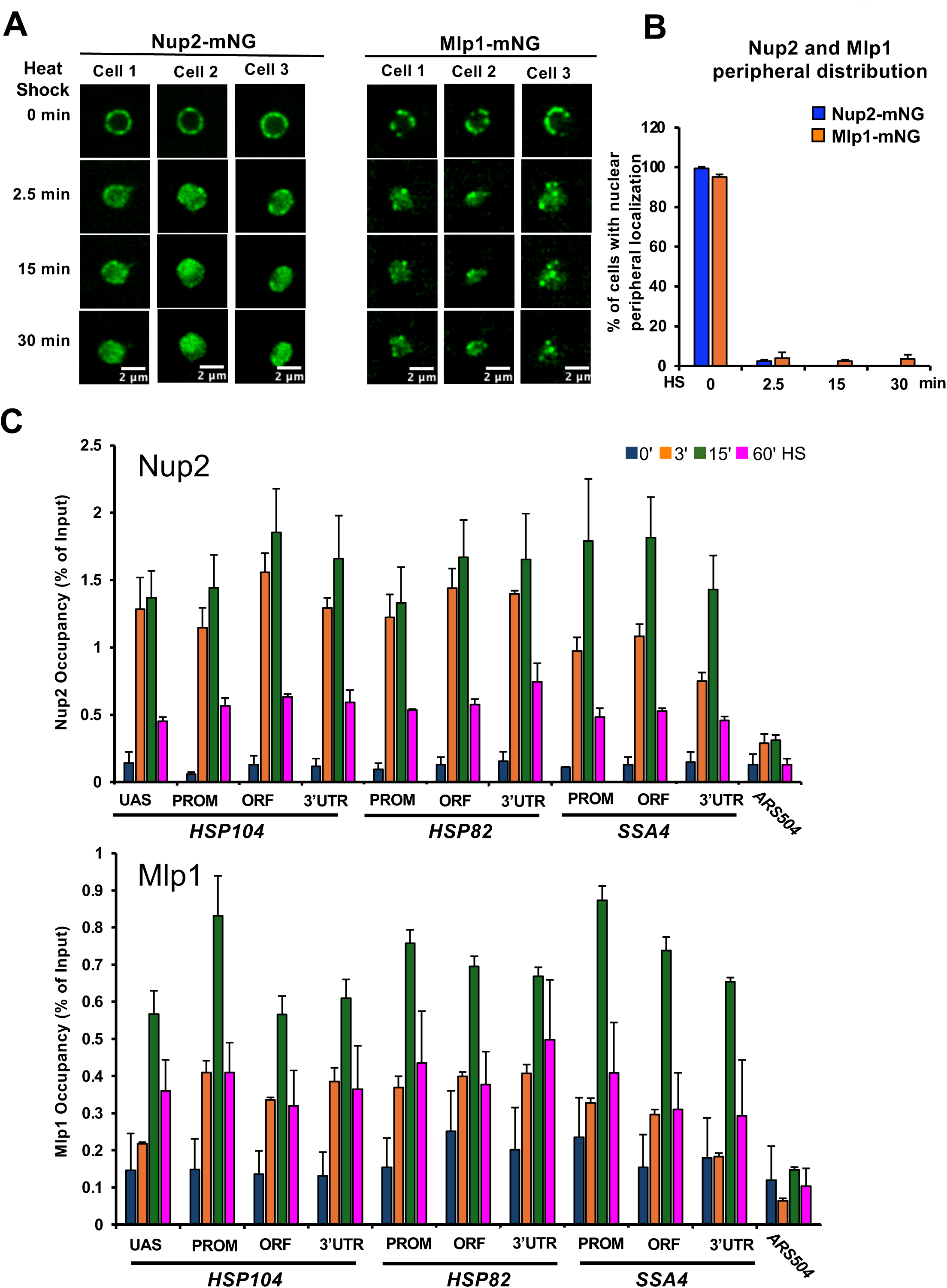
Nup2 and Mlp1 rapidly relocalize from nuclear periphery to nuclear interior in cells exposed to heat shock and are recruited to *HSR* genes with similar kinetics. (A) **Left Panel Set:** Micrographs showing the distribution of Nup2-mNG in SMY216 cells under NHS (25°C; 0 min) and HS (39°C; 2.5 - 30 min) conditions. **Right Panel Set:** Same, except cells expressing Mlp1-mNG (SMY192) were evaluated. In both panels, a representative image of the Z stack is shown for each condition. Step size = 0.5 microns. (B) Quantification of cells with a peripheral distribution of Nup2-mNG or Mlp1-mNG over a HS time course. Approximately 100 cells were scored per biological sample per time point. Depicted values are Means + SD, N=2. (C) Nup2 and Mlp1 occupancy of *HSR* genes is greatly enhanced following heat shock. Occupancy of the indicated loci – UAS, promoter (PROM), transcribed region (ORF) and 3’-UTR – was evaluated using ChIP-qPCR. Nup2-myc9- and Mlp1-myc9-tagged cells (strains SMY164 and SMY166, respectively) were subjected to the indicated heat shock prior to HCHO-mediated crosslinking, chromatin isolation and immunoprecipitation with mAb 9E10 (see Methods). *ARS504* served as a non-transcribed negative control. Depicted are means + SD, N=2, qPCR=4.

Previous studies have suggested that Nup2 and Mlp1 physically associate with inducible genes upon their activation (70, 71) (see Table 1). Given their HS-induced intranuclear redistribution, we reasoned that Nup2 and Mlp1 may associate with Hsf1 target genes in a HS-dependent manner. To address this, we performed a chromatin immunoprecipitation (ChIP) assay over a similar heat shock time course. This analysis revealed enhanced association of both Nup2 and Mlp1 within regulatory and coding regions of *HSP104, HSP82* and *SSA4* in cells exposed to heat shock relative to the control, non-induced state (Figure 6C). Consistent with its dispersed distribution within the nucleoplasm, Nup2 binding to these *HSR* genes was more rapid and of a greater magnitude than that of Mlp1. Notably, the association of each protein was transient, as occupancy in both cases peaked at 15 min and diminished thereafter. Importantly, the association of Nup2 and Mlp1 with a non-transcribed locus (*ARS504*) remained at near-background levels over the entire time course (Figure 6C).

### Nup2 and Mlp1 are dispensable for the formation of heat shock-induced transcriptional condensates

Transcriptional condensates, enriched in Hsf1, Pol II and Mediator, form in response to heat shock and have been implicated in driving the spatial rearrangement of *HSR* genes (18). It has been suggested that there is a functional link between Hsf1 condensate formation and *HSR* intergenic interactions (17–19). As described above, Nup2 and Mlp1 are required for *HSR* gene coalescence (Figures 5F and 5G), and this is likely mediated through heat shock-induced association of Nup2 and Mlp1 with *HSR* genes (Figure 6C). It was therefore of interest to investigate whether Nup2 and Mlp1 are also required for the formation of heat shock-dependent Hsf1 condensates. To address this, we imaged Hsf1-mNG in no degron and double degron (Nup2-, Mlp1-AID) cells over a heat shock time course (experimental strategy summarized in Figure 7A). As previously observed (17–19), Hsf1 forms heat shock-dependent subnuclear clusters in Nup2-, Mlp1-expressing (no degron) cells (Figures 7B and S5A). Despite their role in spatial genome organization, simultaneous depletion of Nup2 and Mlp1 had little or no effect on the fraction of cells exhibiting ≥2 Hsf1-mNG foci (Figure 7C). A similar outcome was evident in single *nup2Δ* and *mlp1Δ* deletion strains (Figure 7E). However, the number of foci per cell was somewhat higher in the double depletion strain, particularly at very early time points in a heat shock time course (Figure 7D). Nonetheless, neither Mlp1 nor Nup2 was enriched within Hsf1 condensates in heat-shocked cells relative to the non-shocked control (Figure S6). Collectively, these results argue that Nup2 and Mlp1 promote *HSR* gene interactions without having a substantial impact on the formation of Hsf1 condensates.

**Figure 7.**
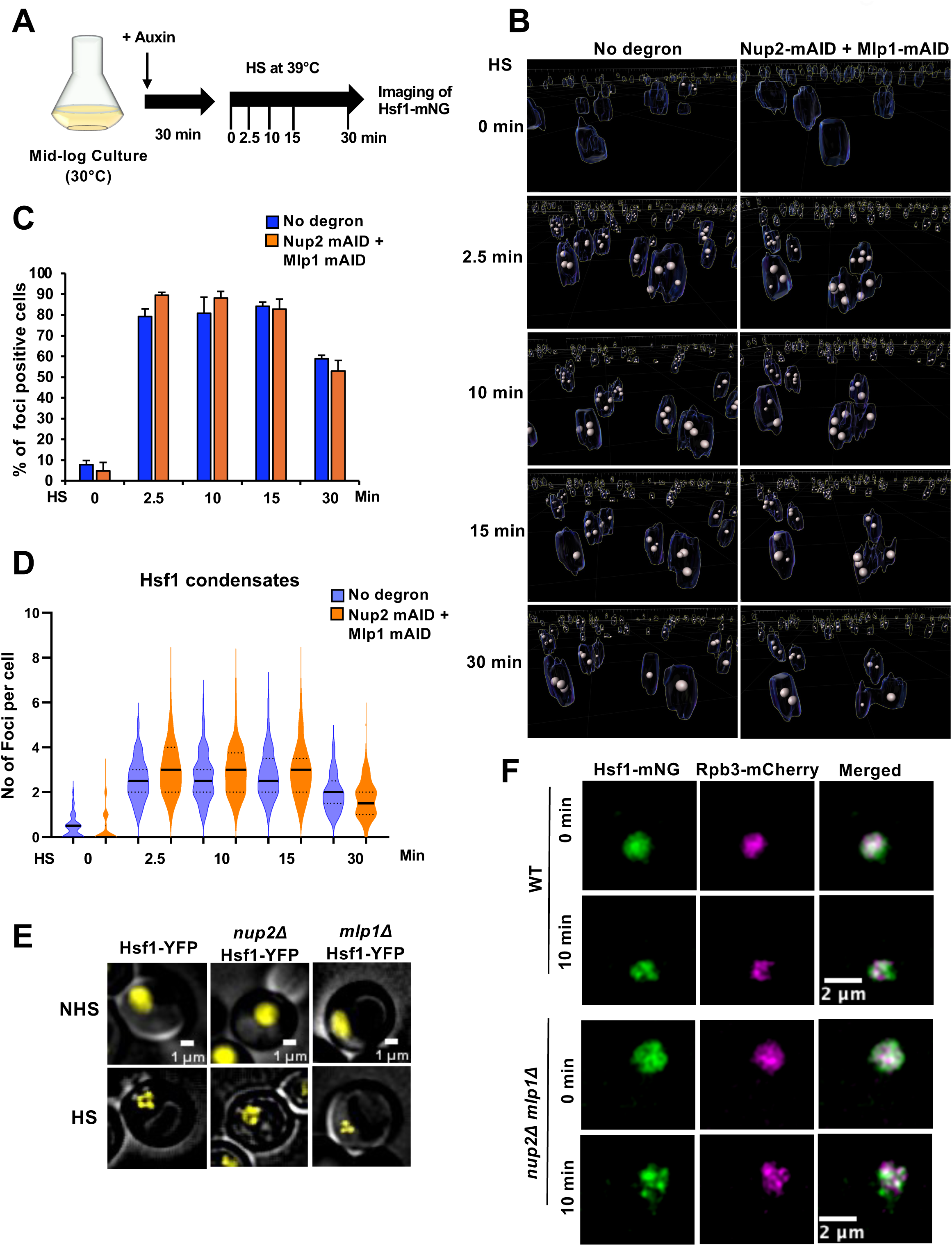
Formation of Hsf1- and Pol II-containing transcriptional condensates is unimpeded in Nup2-, Mlp1-depleted cells. (A) Experimental strategy for live cell imaging. Mid-log phase cultures of Hsf1-mNG cells bearing no degron (SMY172) or the double Nup2-, Mlp1-degron (SMY170) were pretreated with 0.5 mM IAA for 30 min at 30°C, attached to a ConA-coated VAHEAT substrate, subjected to HS for the indicated times and then imaged on a confocal fluorescence microscope. (B) 3D reconstruction of confocal fluorescence micrographs of Hsf1-mNG foci in WT (no degron) and Nup2-, Mlp1-depleted cells over the indicated HS time course. Performed using Imaris v10.1.0 from 11 Z-planes. To enhance separation of Hsf1-mNG foci, the Z-axis was visually stretched relative to the X-Y axes. An outline of each nucleus is shown. Representative deconvoluted images with an accurate scale bar are presented in Figure S5A. (C) Percent SMY172 and SMY170 cells showing ≥2 Hsf1-mNG foci over a HS time course. An average of 150 cells per time point, per biological sample, were quantified (Imaris v.10.1.0). N=2. Values depicted are means + SD. (D) Violin plot showing the distribution of the number of Hsf1 foci per cell. Approximately 300 cells derived from two independent cultures were analyzed per time point for each condition. Black horizontal lines represent the median number of foci per cell. (E) Subnuclear localization of Hsf1-YFP in WT, *nup2Δ* and *mlp1Δ* cells (DPY032, SMY134 and SMY136, respectively) under NHS (25°C) and HS (38°C; 10 min) conditions as indicated. Cells were imaged on a widefield fluorescence microscope. Depicted are representative Z-planes (step size 0.5 microns). (F) Subnuclear localization of Hsf1-mNG and Rpb3-mCherry in WT and *nup2Δ mlp1Δ* cells (LRY040 and LRY120, respectively) under NHS and HS (39°C) conditions as indicated. Cells were imaged using confocal microscopy. Depicted are representative Z-planes (step size 0.5 microns).

Recently, it has been argued that in addition to the formation of Hsf1 condensates, Pol II incorporation into these transcriptional condensates is an obligatory step in driving *HSR* gene coalescence in heat-shocked cells (72). To test whether Pol II condensation is blocked in *nup2Δ mlp1Δ* cells, we constructed a double delete strain expressing both Hsf1-mNG and Rpb3-mCherry and imaged the strain under both NHS and HS conditions as above. As previously observed (18), Rpb3 condenses and a fraction colocalizes with Hsf1 following HS in WT cells (Figure 7F and S5B). Notably, virtually identical condensation and colocalization were observed in the corresponding *nup2Δ mlp1Δ* strain. This result argues that the failure of *HSR* genes to undergo 3D restructuring in double depletion mutants arises from a step downstream of mature transcriptional condensate formation (see Figure 10 below for model).

### Nup2 and Mlp1 are not required for the recruitment of Hsf1 or Pol II to *HSR* genes

Given that heat shock-induced Hsf1 clusters are associated with *HSR* genes (18) and Hsf1 occupancy upstream of these genes increases significantly following a brief HS (16, 73–75), we wished to know whether Nup2 and Mlp1 promote the occupancy of Hsf1 at representative UAS regions. To address this, we simultaneously depleted both proteins as above and performed ChIP following 0, 3 and 15 min of HS. As shown in Figure 8A, Hsf1 occupancy was not affected by the simultaneous depletion of Nup2 and Mlp1, indicating that these proteins play little or no role in the recruitment of Hsf1.

**Figure 8.**
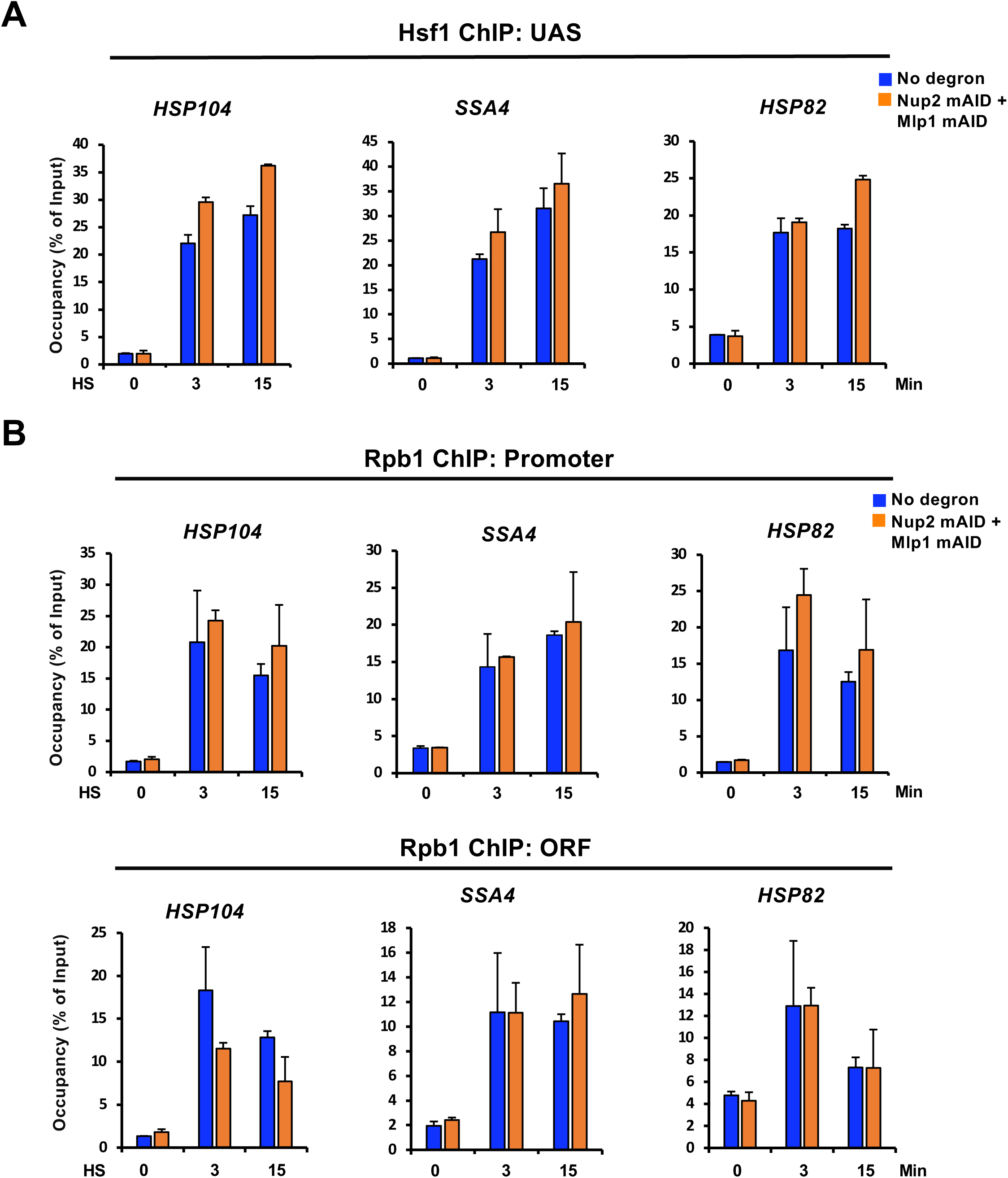
Hsf1 and Pol II occupancy of *HSR* genes is unimpeded in Nup2-, Mlp1-depleted cells. (A) Hsf1 ChIP of the UAS regions of representative *HSR* genes in no degron (LRY016) and double degron (SMY152) strains pretreated with 0.5 mM IAA for 30 min before exposing them to HS for the indicated time points. Values are depicted as means + SD. N=2, qPCR=4 (B) Rpb1 ChIP of the promoter and ORF regions of representative *HSR* genes conducted in control and Nup2-, Mlp1-double depletion strains as in (A).

We next asked whether Nup2 and Mlp1 are involved in recruiting Pol II to the *HSR* genes. We depleted them together and performed ChIP of Rpb1 (Pol II large subunit) as above. ChIP analysis demonstrated that simultaneous depletion of Nup2 and Mlp1 had no detectable effect on Pol II occupancy at either the promoter or coding region of representative *HSR* genes (Figure 8B), indicating that the nuclear basket proteins are not critical for the heat shock-induced recruitment of Pol II to the *HSR* genes. Collectively, these observations suggest that double depletion of Nup2 and Mlp1 suppresses *HSR* gene coalescence in acute heat-shocked cells without diminishing the chromatin association of either Hsf1 or Pol II.

### Nup2 and Mlp1 are not required for transcriptional induction of the *HSR* genes

It has been suggested that TF condensates are a key mechanism for transcriptional regulation (76–80). Previous studies have noted that heat shock triggers strong transcriptional induction of *HSR* genes (16, 58, 81) and there is a temporal correlation between Hsf1 condensation and transcription in yeast cells exposed to acute heat shock (17–19). In the experiments described above, we observed that simultaneous depletion of Nup2 and Mlp1 had no effect on Hsf1 condensate formation. To ask whether simultaneous depletion of these proteins affects the heat shock-induced transcriptional activation of *HSR* genes, we conducted RT-qPCR (Figure 9A).

**Figure 9.**
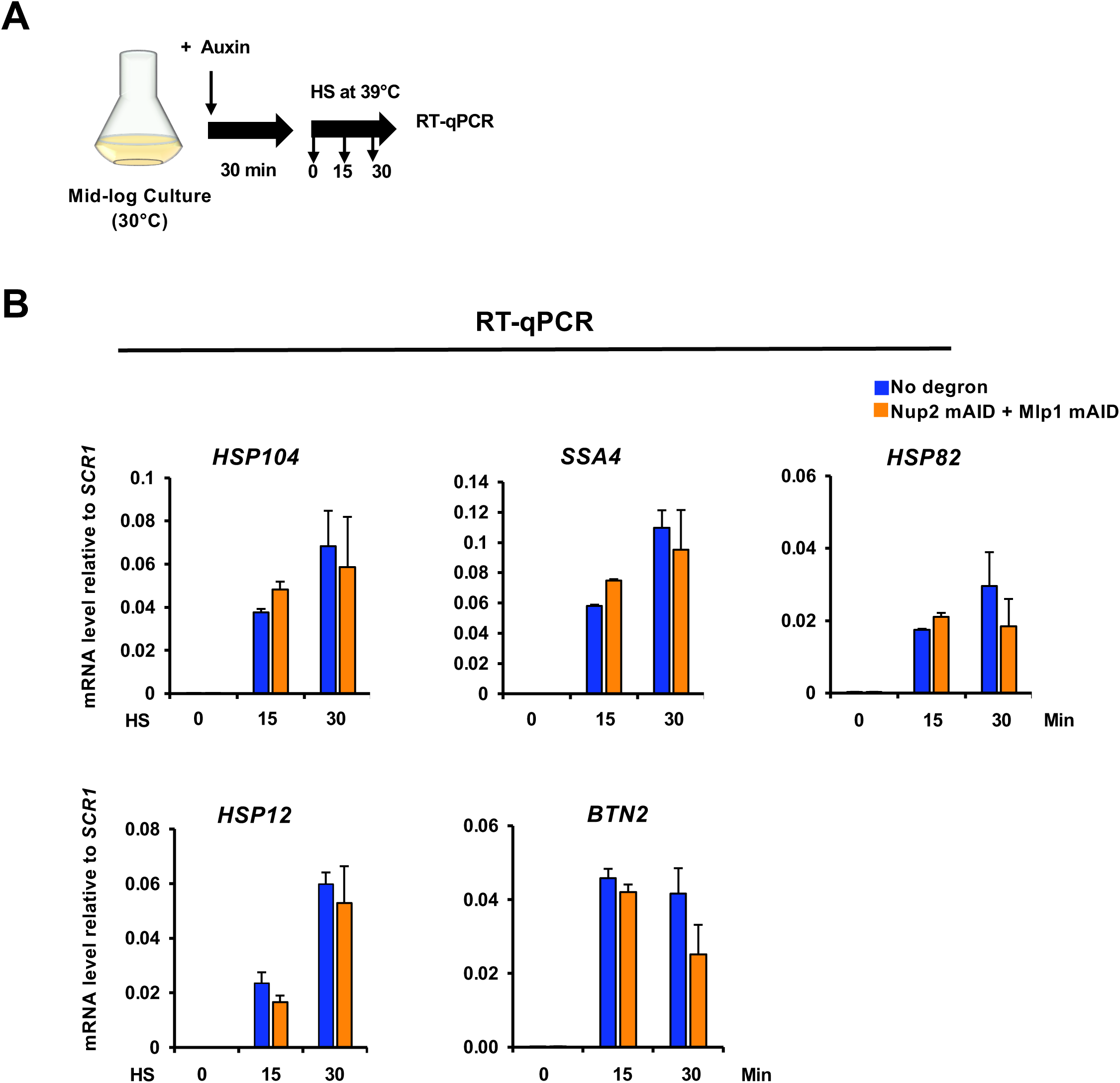
Transcriptional induction of *HSR* genes occurs unimpeded in Nup2-, Mlp1-depleted cells. (A) Experimental strategy for assaying *HSR* transcript abundance. Mid-log phase (A_600_ =0.5) LRY016 (no degron) and SMY152 (Nup2- and Mlp1-double degron) cultures were grown in liquid YPDA and pretreated with 0.5 mM IAA for 30 min prior to exposing them to HS for the indicated durations. Total cell RNA was then isolated, and the indicated mRNAs were quantified by RT-qPCR. (B) *HSR* mRNA levels in LRY016 and SMY152 cells were determined as described above and their abundance normalized to the Pol III transcript *SCR1*. Depicted are means + SD. N=2, qPCR=4.

Consistent with the absence of an effect on Hsf1 condensate formation, we observed that double depletion of Nup2 and Mlp1 failed to impact *HSR* mRNA abundance under either the NHS or HS condition (Figure 9B). This finding is in accord with previous work which found that deletion of *MLP1* has no impact on the transcriptional activation of either *GAL10* or *HSP104* genes (66). It is also consistent with a recent study reporting that depletion of Nup2 does not affect genome-wide mRNA levels (82).

## Discussion

We have provided new insights into a signature feature of the heat shock transcriptional response – *HSR* gene coalescence (HGC) – and the factors that contribute to the spatial repositioning of *HSR* genes. It has been previously observed that *HSR* genes can relocate to the NPC following their transcriptional activation (22, 66). Consistent with these earlier observations, we detect coalescence of *HSR* genes at the nuclear periphery. More frequently, however, such coalescence occurs within the nucleoplasm. Moreover, using complementary imaging and molecular approaches, we have shown that the nuclear basket proteins Nup2 and Mlp1, inducibly recruited to *HSR* genes, are required to drive these genes into coalesced chromatin clusters. Previous studies suggested that inactivation of either Mlp1 or Nup2 obviated peripheral localization of transcriptionally induced *HSR* genes (22, 66). However, we observed that the deletion of either Mlp1 or Nup2 typically had only a mild effect on *HSR* gene clustering. This suggests that the two basket proteins play functionally redundant roles in regulating yeast 3D genome organization. Interestingly, association of *Drosophila* Nup50 (orthologue of Nup2) with heat shock chromosomal puffs has been previously observed (83), suggesting that association of nuclear basket proteins with activated *HSR* chromatin is evolutionarily conserved.

In contrast to Mlp1 and Nup2, we found that two essential, scaffold-associated proteins, Nup1 and Nup145, play no detectable role in driving intergenic interactions between *HSR* genes in heat shock-induced cells. A previous study suggested that Nup1 is involved in subnuclear repositioning and interallelic clustering of *GAL1-10* genes (57), yet we observed that depletion of Nup1 (in combination with Nup145) had minimal impact on *HSR* gene coalescence.

Our findings contrast with the clustering of *MET* genes that occurs during methionine starvation (23, 84), as well as with the interallelic clustering of galactose-inducible and inositol starvation-inducible genes that occurs upon their transcriptional activation (57). In these cases, the clustering appears to occur at the NPC and NPC tethering promotes the transcription of these genes. In notable contrast, we show that *HSR* gene clustering occurs downstream of *HSR* transcriptional induction. Furthermore, as mentioned above, we show that *HSR* gene clustering occurs predominantly in the nucleoplasm and this fact, in combination with the heat shock-induced subnuclear relocalization of Nup2 and Mlp1, argues that the nuclear basket proteins mediate their topological effects in an untethered, NPC-free state.

A second signature of the heat shock transcriptional response is the inducible formation of Hsf1 condensates that colocalize with *HSR* genes (18), a phenomenon also observed in human cells (85). In budding yeast, Hsf1 condensates form rapidly in response to acute HS (detectable within 2.5 min of a 30° to 39°C shift) but begin to return to a diffuse state as early as 30 min (this study and (17–19). These condensates have been functionally linked to the 3D repositioning of *HSR* genes (18). Several factors, including Hsf1, Pol II and Mediator, have been implicated in driving both Hsf1 condensate formation and *HSR* gene coalescence^1^ (17–19). Nup2 and Mlp1 appear to represent a distinct category of nuclear factors, since they are required for *HSR* gene clustering yet unlike the aforementioned factors are dispensable for the formation of Hsf1 condensates and indeed do not colocalize with Hsf1 in HS-induced cells. They also play no detectable role in the recruitment of either Hsf1 or Pol II to *HSR* genes. These results indicate that in response to HS, the formation of Hsf1 condensates and recruitment of Hsf1 and Pol II to *HSR* genes can occur in the absence of long-range topological restructuring (see Figure 10 for model).

**Figure 10.**
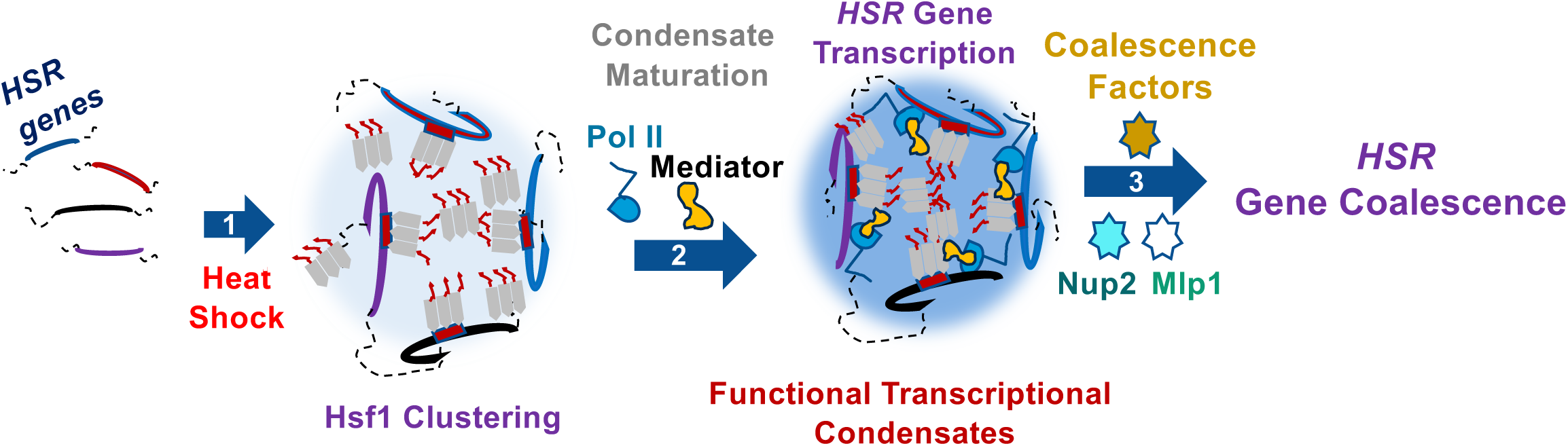
Model: Nup2 and Mlp1 promote *HSR* gene coalescence without being incorporated into transcriptional condensates. **1.** In response to heat shock, Hsf1 trimerizes and binds to the HSEs upstream of *HSR* genes (Hsf1 clustering). Some Hsf1 clusters may exist as clouds of non-DNA-bound molecules (19, 115). **2.** RNA Pol II and Mediator are recruited to the Hsf1 cluster and Hsf1 condensates undergo maturation (18, 95). This functional condensate initiates *HSR* gene transcription. **3.** Following formation of Hsf1 condensates and transcriptional activation of Hsf1 target genes, Nup2 and Mlp1 are recruited to *HSR* genes but not detectably incorporated into Hsf1 condensates. This physical association with chromatin induces *HSR* gene clustering. The coalesced gene foci are predominantly detected within the nucleoplasm.

How might Nup2 and Mlp1 elicit their topological effect? Nup2 is involved in nucleocytoplasmic transport of proteins and RNA (39, 86) while Mlp1 binds RNA export factors and participates in mRNA quality control and export (67, 87, 88). Results presented here are compatible with these primary roles as we have shown that in heat shock-induced cells, Nup2 and Mlp1 [i] do not colocalize with heat shock-induced Hsf1 (as detected by live cell microscopy); [ii] do not impact either Hsf1 or RNA Pol II recruitment to *HSR* genes; and [iii] do not impact total *HSR* mRNA abundance. Yet both proteins undergo marked intranuclear relocalization upon HS and both physically associate with *HSR* genes. Given the predominant diffuse nucleoplasmic distribution of Nup2 and its strong occupancy of *HSR* genes, evident as early as 3 min following temperature upshift, Nup2 may directly drive *HSR* gene clustering via its ability to dynamically exchange between the nucleoplasm and NPC (36, 39). Although Mlp1 might contribute to this shuttle-and-clustering activity in heat-shocked cells given its relocalization from the nuclear periphery, diffuse distribution within the nucleoplasm and inducible occupancy of *HSR* genes, it is notable that a fraction of Mlp1 molecules condense into several discrete foci upon HS (Figures 6A, 6B). This latter phenomenon has been observed by others (68, 69). Therefore, diffusely distributed Mlp1 may directly promote the 3D spatial repositioning and clustering of *HSR* genes, while the foci-associated fraction of Mlp1 may serve an indirect regulatory role by sequestering inhibitory factors of gene coalescence. Of relevance, heat shock-dependent sequestration of mRNA export factors Nab2 and Yra1 was observed in Mlp1 foci (69). This sequestration role may explain why we see a strong HGC phenotype despite the presence of Mlp2 (paralogue of Mlp1): either Mlp2 does not participate in the formation of such condensates or Mlp1/Mlp2 condensation may require a threshold concentration of the two proteins.

Consistent with our observations of Nup2 and Mlp1 enhancing both E-P looping and intragenic folding, Nup2 has previously been reported to exhibit boundary/insulator properties (89, 90) and Mlp1 has been implicated in gene looping (91). Interestingly, none of the intergenic or intragenic perturbations arising from Nup2-, Mlp1-double depletion materially impacts the total abundance of *HSR* mRNA following acute exposure to HS. These observations suggest that physical proximity between regulatory elements as detected by 3C analysis is unnecessary for robust Hsf1-driven transcription of representative genes. This is consistent with recent models of mammalian and *Drosophila* transcription that argue against the necessity of physical contact between enhancer and promoter elements to instigate transcriptional induction (92–94). Collectively, our findings are consistent with a recent model proposing that transcription factor clustering and the formation of transcriptional condensates may occur prior to the spatial coalescence of enhancers or the formation of transcriptional hubs (95).

### Conclusions and limitations of this study

In conclusion, our observations reveal a novel, highly specific role for the nuclear basket proteins Nup2 and Mlp1 in promoting the 3D repositioning and coalescence of heat shock-induced *HSR* genes. This role is likely served when these proteins are in their NPC-free, diffusive state. Other phenomena associated with the heat shock transcriptional response – Hsf1 condensation, Hsf1 binding to upstream regulatory regions of *HSR* genes, Pol II condensation, Pol II recruitment and transcription of *HSR* genes – occur independently of Nup2 and Mlp1. In this regard, our work has identified a unique category of factors, since other factors thus far characterized in the *HSR* lack such specificity and appear to participate in most if not all steps (17–19).

The findings presented here provide further evidence that Hsf1 condensate formation can be uncoupled from downstream events in the heat shock response. We recently showed that in response to 8.5% ethanol stress, Hsf1 condensates form and *HSR* genes coalesce well before *HSR* genes are transcriptionally activated (19). Here, we have demonstrated that in response to thermal stress, Hsf1 condensate formation can occur without downstream *HSR* gene repositioning in Nup2-, Mlp1-depleted cells. While the biological consequence of aberrant 3D *HSR* gene organization in *nup2 mlp1* mutants remains unclear, recent evidence suggests that HGC promotes *HSR* RNA multiplexing (96), a phenomenon that could contribute to the coordinate regulation of *HSR* genes and proper subcellular colocalization of their encoded mRNAs (97).

Limitations of this study include the following. We do not know the mechanism by which Nup2 and Mlp1 reshape 3D genome architecture, including the identity of molecular players downstream of the NB proteins. We do not know the mechanism by which Nup2 and Mlp1 are recruited to *HSR* gene regulatory and coding regions (or for that matter, whether their role is different when associated with regulatory versus coding regions). We have not established whether Nup2 and Mlp1 redistribution following HS is reversible in heat-shocked cells. We note, however, that such reversal is unnecessary for either dissolution of Hsf1 condensates or dissipation of HSR intergenic interactions, as both phenomena appear within the first 30 min of HS (Figure 7; (18, 19, 58)). Finally, it remains unclear what the biological significance of Hsf1 condensate formation and *HSR* gene coalescence is. It is possible that *HSR* gene clustering promotes *HSR* mRNA multiplexing as mentioned above. Other possibilities, including enhancement of post-transcriptional processing and nucleocytoplasmic export of *HSR* mRNA, are currently under investigation.

## Experimental Procedures

### Yeast Strain Construction

For microscopy analyses, to construct SMY206, we initially PCR-amplified the *KAN-MX* gene from pFA6a-KanMX6 using forward and reverse chimeric primers containing ∼50 bp of *HSP82* after the poly (A) site juxtaposed against 20 bp sequence homologous to the plasmid. We transformed ASK726 with the PCR amplicon to insert *KAN-MX* downstream of the *HSP82* gene, creating SMY108. *KAN-MX* served as a landing pad to insert *TetO_200_::LEU2*. (Note: actual TetO repeat length is likely <200 bp.) The plasmid pSR14 (*TetO_200_::LEU2),* kindly provided by S. Gasser, Friedrich Miescher Institute for Biomedical Research, was linearized with Asc I, creating homologous ends for *KAN-MX*. The linearized *TetO_200_::LEU2* construct was inserted into the *KAN-MX* locus using allelic replacement, creating SMY109. SMY118 was created by crossing SMY109 with W303-1B, followed by sporulation, tetrad dissection, and selection of a haploid bearing only the *LEU2* marker. SMY110 was created by crossing ASK701 with W303-1A followed by sporulation, tetrad dissection, and selection of a haploid with only the *TRP1* marker. Plasmid pMY63 (*REV1pr-tetR-mCherry*, *REV1pr-lacI-GFP*), a gift of Lu Bai, Penn State University, was then linearized by digestion with BsiWI and inserted into the *his3* locus of SMY110, creating SMY123. The SMY206 diploid was generated by crossing SMY123 with SMY118.

Strain SMY207 was constructed by crossing SMY123 and SMY118 following deletion of *MLP1* from both strains using the *mlp1Δ::KAN-MX* PCR product as transforming DNA (genomic template obtained from the ResGen KO strain). SMY208 was generated in a similar fashion using *nup2Δ::KAN-MX* as the transforming DNA. SMY201 was constructed by crossing ASK722 x ASK726 following deletion of *NUP2* from both strains using *nup2Δ::KAN-MX*. SMY203 was generated in a similar fashion using *mlp1Δ::KAN-MX*. SMY134 and SMY136 were generated by deleting *NUP2* and *MLP1*, respectively, from DPY032. SMY160, SMY163, and SMY182 were created by transforming LRY016, SMY148, and SMY152, respectively, with *POM34-mCherry::NAT* using JTY001 genomic DNA as a template. SMY170 and SMY172 were constructed by transforming SMY152 and LRY016, respectively, with *HSF1-mNeonGreen::HIS5* using LRY037 genomic DNA as a template. SMY192 and SMY196 were constructed by in-frame insertion of *mNeonGreen::HIS5* at the C-terminus of *MLP1* in LRY016 and SMY148, respectively. SMY216 and SMY221 were constructed in a similar fashion by targeting the C-terminus of *NUP2* in LRY016 and SMY148. The plasmid template for these amplifications was pFA6a-link-ymNeonGreen-SpHis5 (ymNeonGreen = yeast optimized mNeonGreen). LRY777 and LRY888 were constructed by transforming SMY192 and SMY216, respectively, with the *HSF1*-targeted *mCherry::URA3* amplicon. LRY119 was generated by crossing strains LRY117 and LRY118, both of which had deletions of *NUP2* and *MLP1* using *nup2Δ::KAN-MX* and *LoxP-klURA3-Lox* cassette flanked by 50 bp of 5’- and 3’-regions of *MLP1* outside of ORF for homologous recombination as the transforming DNA. *LoxP-KlURA3-LoxP* cassette was obtained from plasmid pUG72 by PCR amplification. LRY120 was derived from LRY040, with *NUP2* and *MLP1* deleted using the same method described above.

For molecular analyses, SMY143 and SMY145 were constructed by introducing a mini-degron tag amplified from *pKAN-mAID*-9myc* (98) into LRY016, respectively, targeting the C-termini of *NUP1* and *MLP1*. Similarly, strains SMY148, SMY149, and SMY152 were generated by introducing a mini-degron tag PCR amplified from *pHyg-mAID*-9myc* (98) into the C-terminus of *NUP145* in SMY143, *NUP2* in LRY016, and *NUP2* in SMY145, respectively. SMY164 and SMY166 were constructed by in-frame insertion of a *Myc9::TRP1* tag at the C terminus of *NUP2* and *MLP1* in strain LRY016 using pWZV87 as PCR template.

See Tables S1, S2, and S3 for complete lists of strains, plasmids, and primers used in strain construction.

### Yeast Culture and Treatment Conditions

Yeast cells were grown overnight (O/N) in liquid YPDA (1% w/v yeast extract, 2% w/v peptone, 2% w/v dextrose, and 20 mg/L adenine). Cells from the overnight cultures were inoculated into fresh liquid YPDA and allowed to grow at 30°C with constant shaking until they reached mid-log density (OD_600_ = 0.5 to 0.8). Then the cultures were incubated at 25°C for 10 min prior to heat shock. To induce heat shock, the mid-log phase culture was combined with an equal volume of YPDA medium preheated to 55°C to elicit an instantaneous 25°C to 39°C shift. The culture was then maintained at 39°C for the specified duration. Control samples without heat stress were diluted with an equal volume of YPDA and kept at 25°C. All samples were maintained at their respective temperatures using a water bath with continuous shaking.

### Auxin-Induced Degradation (AID) and Immunoblot Analysis

The proteins of interest used in this study were degraded using the AID strategy (98) To optimize auxin concentration and incubation time, actively growing (OD_600_=0.5) degron-tagged cells expressing the F-box protein osTIR1 were treated with different concentrations of indole-3-acetic acid (0.5 or 1 mM IAA) at 30°C for 0, 20, 30 and 60 min before undergoing metabolic arrest with 20 mM sodium azide, followed by cell harvesting. Fresh IAA stocks (100 mg/mL [570 mM]) were prepared in 95% ethanol and sterilized by filtration before use. For the control (0 min) sample, cells were treated with an equal volume of vehicle alone.

The harvested cells were subjected to protein extraction and immunoblot analysis as previously described (65). Signal intensities of protein bands were quantified using Image Lab software (version 5.2.1, Bio-Rad) and normalized to the loading control Pgk1. Monoclonal antibodies (mAbs) targeting Myc (9E10; Santa Cruz Biotechnology sc-40) and Pgk1 (ThermoFisher 459250) were used for detection.

For 3C, ChIP, RT-qPCR and microscopic analysis, mid-log cells grown in YPDA or synthetic dextrose complete (SDC) medium were treated with 1 mM IAA for 60 min (for LRY016, SMY148, SMY160 and SMY163) or 0.5 mM IAA for 30 min (for LRY016, SMY145, SMY149, SMY152, SMY160, SMY170, SMY172 and SMY182) prior to +/- heat shock for the specified time points. Auxin concentrations were kept constant throughout the experiment (+/- HS) to ensure continuous degradation of the target proteins.

### Spot Dilution Assay

O/N cultures grown in liquid YPDA at 30°C were diluted to OD_600_=0.5 using sterile distilled water. Cells were then serially diluted 1:5 and 7 μl of each dilution were spotted on YPDA plates using a 20 μl pipette. The plates were incubated for 36 to 48 hours at 24°C, 30°C, 35°C, and 37°C (Figure S2).

### Cell Viability Assay

Mid-log cells (OD_600_=0.5), obtained following inoculation of fresh liquid YPDA medium with an O/N culture, were exposed to heat shock (39°C) for the indicated time points (Fig. S2) using the method described above. Cultures were diluted 1: 10,000 and plated onto YPDA plates. Plates were incubated at 30°C for 36 to 48 hours. Colonies were counted and the average number from two replicates was plotted on a bar graph. For the Nup2 and Mlp1-double degron-tagged strain (SMY152), mid-log cells (OD_600_ 0.5), obtained following inoculation of fresh liquid YPDA medium with an O/N culture, were treated with 0.5 mM auxin (IAA) or vehicle alone for 30 min at 30°C. These cells were then subjected to +/-heat shock, as mentioned above. Auxin concentration was maintained throughout and cells from each heat shock time point were collected, diluted, and plated onto YPDA plates.

### Taq I - Chromosome Conformation Capture (Taq I-3C)

Taq I-3C (64) was performed essentially as described (65). From O/N cultures, a master cell culture was inoculated and grown in liquid YPDA at 30°C, starting from an initial OD_600_ of 0.15 and reaching a final OD_600_ of 0.65 to 0.7 before subjecting the cultures to +/- HS. For each experimental condition (NHS and HS), 50 mL aliquots of cultures were used. Heat shock was performed as described above. Target proteins were degraded using the AID strategy outlined above before exposing the cells to heat shock. Primers used for analyzing the 3C templates are listed in Supplemental Table S6.

### Reverse Transcription - Quantitative PCR (RT-qPCR)

RT-qPCR was performed as previously described (65). PCR primers used for this analysis are listed in Supplemental Table S4. Target proteins were conditionally degraded as described above before exposing the cells to heat shock.

### Chromatin Immunoprecipitation (ChIP)

Hsf1 and Pol II ChIP assays were performed as previously described (19) using rabbit antisera raised in our laboratory (99, 100). For Nup2 and Mlp1 ChIP, sonicated chromatin lysates were incubated with 2.5 µL of anti-Myc antibody, and the antibody-bound chromatin fragments were captured on Protein G-Sepharose beads (GE Healthcare) by incubating for 16 h at 4°C. All other steps were conducted as previously described (19). Primers used in the ChIP analysis are listed in Supplemental Table S5.

### Fluorescence Microscopy

#### Widefield Microscopy

Live cell imaging of NHS and HS states (Figures 1C, 2F, 4B, 4C, 5E and 7E) was performed essentially as described (19) using an AX70 Olympus epifluorescence microscope equipped with an Olympus Ach 100/1.25-NA objective. Briefly, mid-log phase (A_600_ =0.6) cells grown in synthetic dextrose complete (SDC) medium, inoculated from an O/N culture, were incubated on a concanavalin A (Con A)-coated coverslip for 20 min. Media and unattached cells were removed, followed by addition of fresh media. Cells were and then subjected to instantaneous heat shock (25°C to 38°C) for the indicated time points. For Figures 2F and 5E, cells were pretreated with the specified concentration of auxin (see AID strategy) prior to attachment on the Con A-coated coverslip, followed by imaging at 25°C. Images were captured across nine z planes (Figure 4B and C), 11 z planes (Figures 1C, 2F, 5E) or 4 to 6 z planes (Figure 7E) with a step size of 0.5 microns. Typically, ∼100 cells were counted per biological sample per time point. Note that in Figures 4 and 7E, Hsf1-mYFP foci were processed using the smooth and sharpen image processing functions in FIJI/ImageJ.

#### Spinning Disk Confocal Microscopy

Live cell imaging of mid-log cultures in Figures 2D, 6A, 7B, 7F, S5 and S6 was performed on an Olympus CSU W1 Spinning Disk Confocal System equipped with an UPlan Apo 100x/1.50 NA objective coupled to a sCMOS camera controlled by cellSens Dimension software as described (19). In Figure 2D, mid-log cells were pretreated with vehicle alone or auxin prior to attachment on a Con A-coated VAHEAT (Interherence Gmbh) substrate, followed by imaging at 25°C. In all other figures mid-log cells were treated similarly (where appropriate) before attaching them, followed by instantaneous heat shock (25°C to 39°C) for the indicated times. Images were captured across 21 z planes with a step size of 0.3 µm for Figure 2D, 11 z planes with a step size of 0.5 µm for Figures 6A, 7F, S5B and S6, and 10 z planes with a step size of 0.56 µm for Figure 7B and S5A. ∽100 cells were counted per biological sample per time point.

Image reconstruction and analysis were done using FIJI/ImageJ (v. 1.53t) (101). NPC integrity was determined by examining subcellular localization of the Pom34-mCherry signal and the percentage of cells exhibiting Pom34-mCherry punctate structure (Figure 2E) was counted manually. Quantification of cells with Hsf1-mNeonGreen foci (Figure 7C and D) was performed using the “Cells” feature in Imaris v.10.1.0. Cells with 2 or more foci were included in the Hsf1-mNG foci analysis (Figure 7C). The diameters of the nucleus and Hsf1 foci were set to 2 µm and 0.48 µm, respectively, in Imaris. For the analysis of Hsf1 foci-containing cells, the nuclear volume ranged from 0.8 to 4 µm^3^. For Figures 7B, 7F, S5 and S6, image deconvolution was performed using the Weiner plugin in CellSens Dimension software. In Figure 4, the quantification of cells with *HSR* gene coalescence was accomplished by manually counting the number of cells displaying overlapping green and red dots.

### Statistical Tests

The statistical significance of the differences in mean values for various assays was determined using Microsoft Excel, as specified in the figure legends. A two-tailed, unpaired “t” test with equal variance was performed in each case.

## Data Availability

All data are presented in the main text or Supplemental Figures and Tables. Source/raw data are available upon reasonable request from the corresponding author.

## Acknowledgments

We thank David Pincus, Kelly Tatchell, Arrigo De Benedetti, Nancy Leidenheimer, Jason M. Bodily, Surabhi Chowdhary and Gurranna Male for insightful discussions and technical advice. We thank Amoldeep Kainth for strain construction and Paula Polk and Malgorzata Bienkowska-Haba for assistance in the LSUHSC-Shreveport Research Core Facility (RRID:SCR_024775). We are grateful to Chabbi Govind (Oakland University); Helle D. Ulrich (Institute of Molecular Biology gGmbH); Susan Gasser (Friedrich Miescher Institute for Biomedical Research); Jurg Bahler (University College London); John Pringle (Stanford University School of Medicine); Bas Teusink (Vrije Universiteit Amsterdam); Kim Nasmyth (Research Institute of Molecular Pathology); and Lu Bai (Penn State University) for providing plasmids and reagents. We also appreciate the generosity of David Pincus, Kelly Tatchell, Amoldeep Kainth and Donna and Jason Brickner for providing yeast strains. This work was supported by NIH grants R15 GM128065 and R01 GM138988 awarded to D.S.G. and Ike Muslow predoctoral fellowships awarded to S.M. and L.S.R.

## Footnotes

1 Male, G., Rubio, L.S. and Gross, D.S. 2025. Unpublished observations.

2 A.S Kainth, personal communication.

