## Supplemental Figures for "Nuclear basket proteins Nup2 and Mlp1 drive heat shock-induced 3D genome restructuring downstream of transcriptional activation"

Figure S1

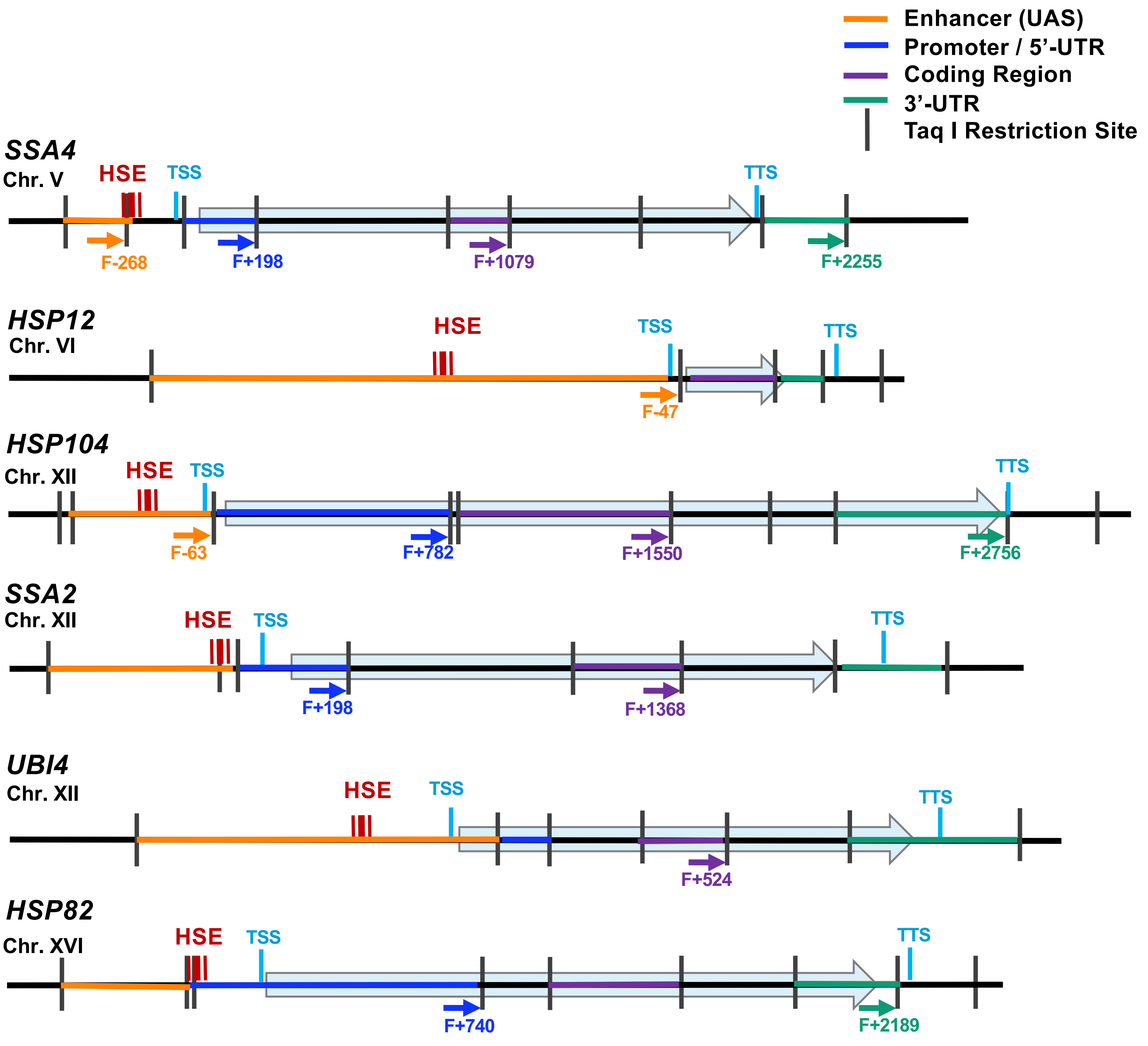


**Figure S1. Location of primers used in 3C analysis and physical maps of Hsf1-regulated genes evaluated in this study.**

Coordinates correspond to Taq I restriction sites (shown as vertical black bars); numbering is relative to ATG (+1). Triple vertical lines (red color) symbolize Hsf1 binding sites (HSEs). TSS, transcription start site. TTS, transcription termination site.

Figure S2

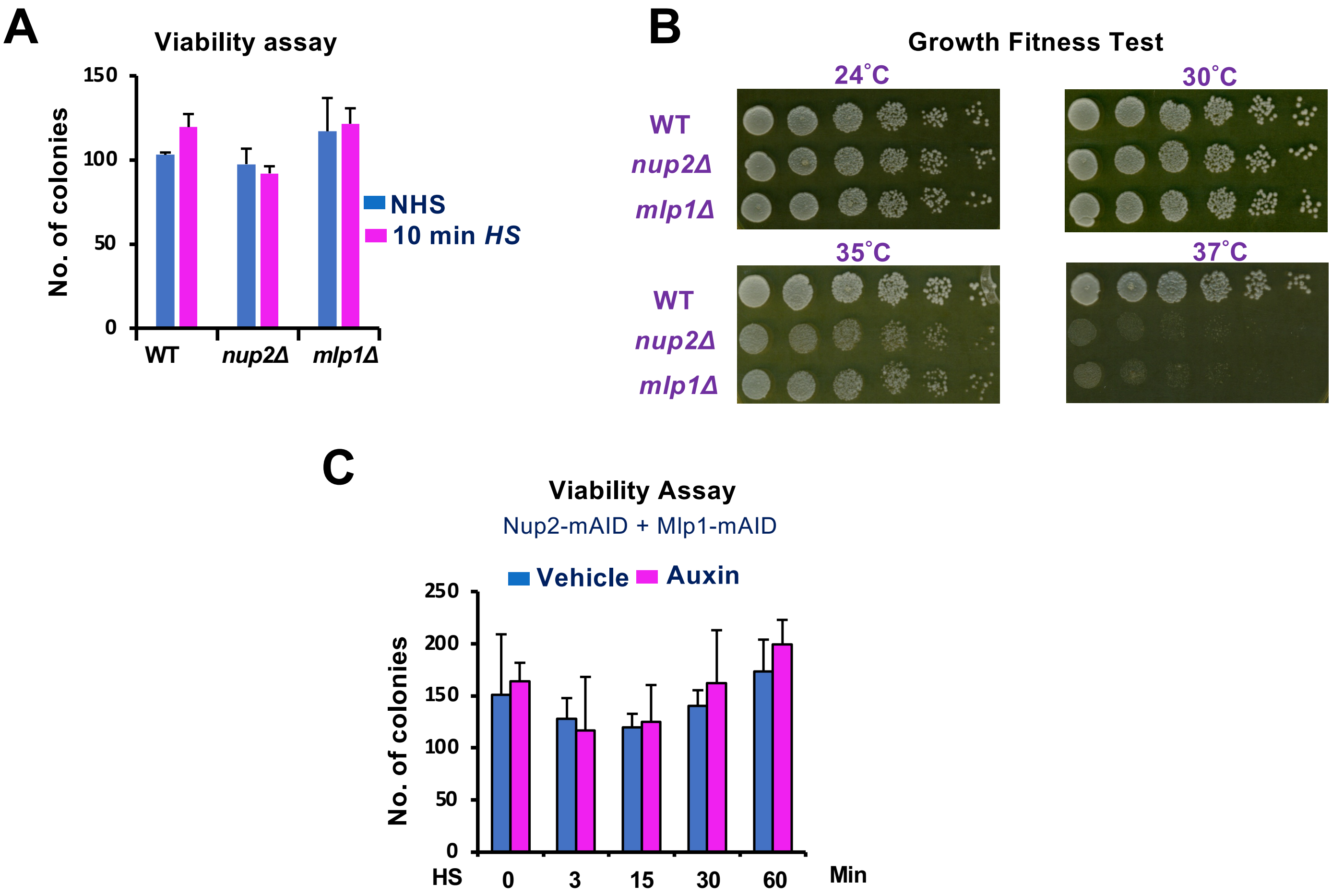


**Figure S2. Viability and Growth Fitness Assays.**

**(A & B) Deletion of either *NUP2* or *MLP1* has no detectable effect on cell viability following brief exposure to 39°C yet diminishes fitness at 37°C.**

(A) Viability assay of strains SMY206 (WT), SMY208 (*nup2Δ/nup2Δ*) and SMY207 (*mlp1Δ/mlp1Δ)* grown on YPDA at 30°C for 1.5 to 2 days following a 10 min heat shock (25°C to 39°C) of a mid-log culture ($A_{600}$ = 0.5). NHS samples were maintained at 25°C. Cultures were diluted 1:10,000 and 100 µl were spread on each plate. The number of visible colonies were counted and depicted as means + SD. N=2.

(B) Spot dilution assay of strains ASK727, SMY201 and SMY203. Five-fold serial dilutions of mid-log cultures were spotted on YPDA plates and incubated at the specified temperatures for 36 to 48 h.

**(C) Double depletion of Nup2 and Mlp1 likewise has no effect on cell viability following brief exposure to 39°C.**

Cell viability of SMY152 (Nup2-mAID + Mlp1-mAID) pretreated with vehicle (0.087% ethanol) or auxin (0.5 mM IAA for 30 min) prior to subjecting cells to HS (39°C) was determined for the indicated times. Cultures were then diluted and spread onto YPDA plates as above and incubated at 30°C for 37 h. The number of visible colonies were counted and depicted as means + SD. N=2.

Figure S3

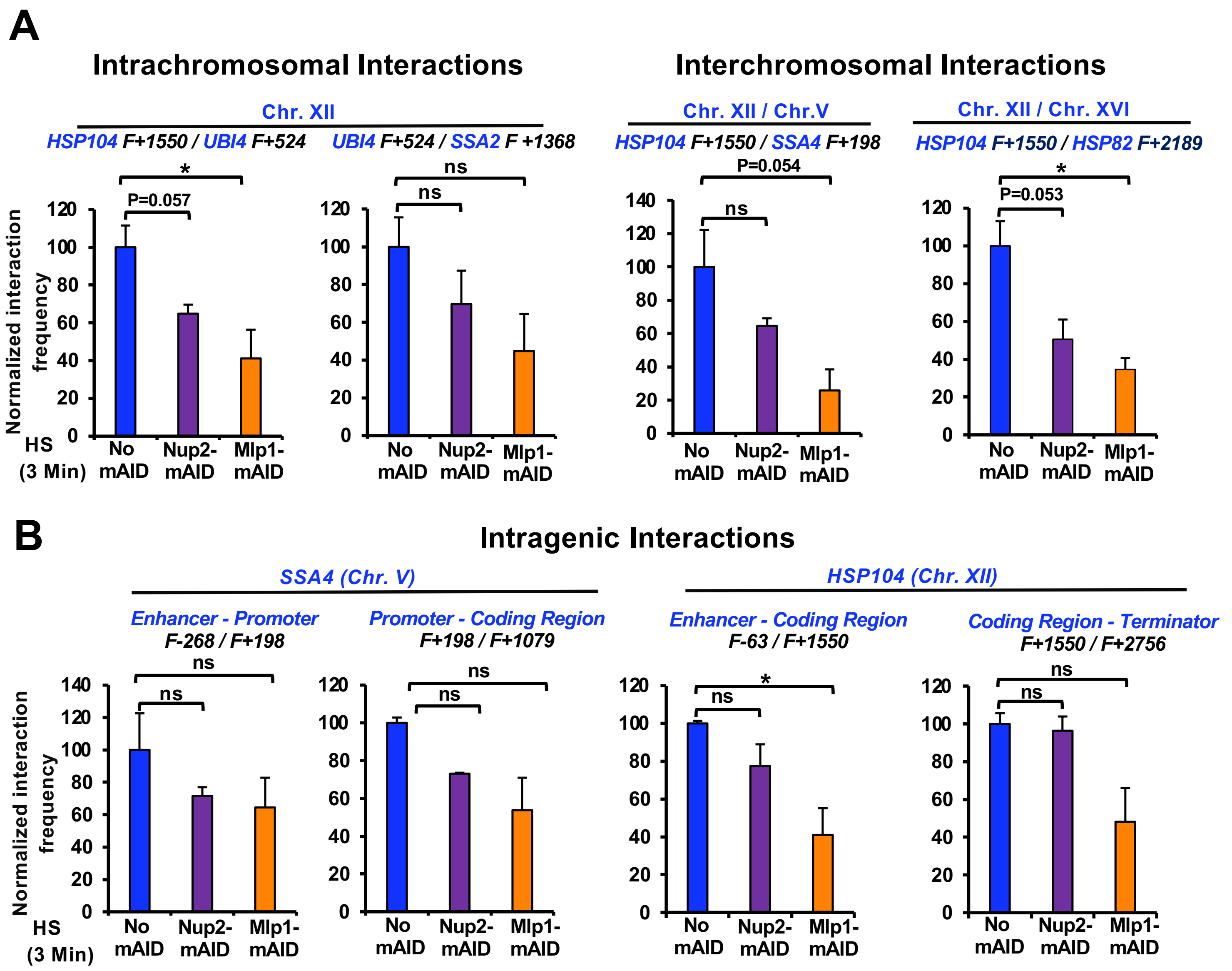


**Figure S3. Acute depletion of either Nup2 or Mlp1 has only a mild effect on *HSR* gene interactions in response to heat shock.**

(A) Taq I-3C assay showing intra- and inter-chromosomal interactions between *HSR* genes in no degron (LRY016), Nup2-mAID (SMY149) and Mlp1-mAID (SMY145) strains pretreated with 0.5 mM auxin for 30 min before exposing them to HS (39°C) for 3 min. Displayed are normalized frequencies of interaction (means + SD; N=2; qPCR=4) as in Figures 3 and 5. Statistical significance was determined as above. *, *P*<0.05; ns, not significant.

(B) Intragenic interactions detected within the *SSA4* and *HSP104* loci determined as above.

Figure S4

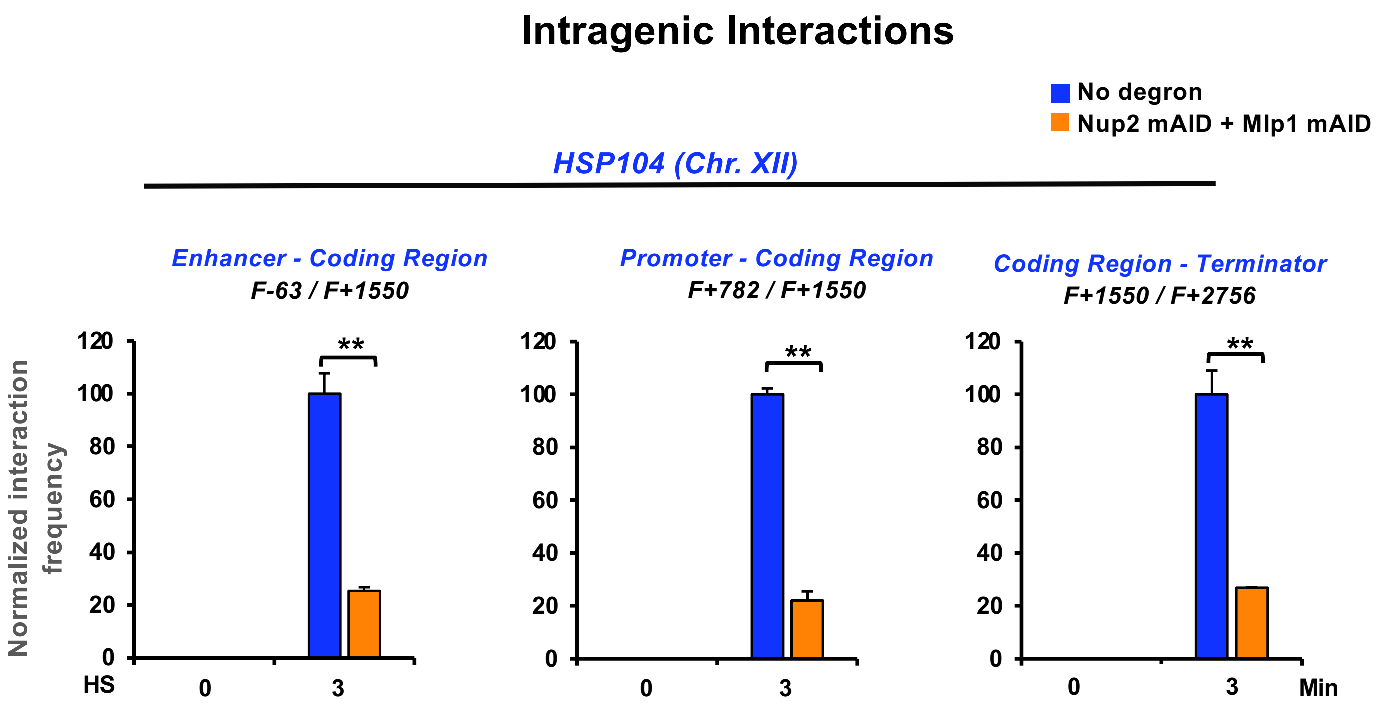


**Figure S4. Nup2-, Mlp1-double depletion significantly reduces frequency of intragenic looping within *HSP104* in acutely heat-shocked cells.**

Intragenic interactions detected within the *HSP104* locus were analyzed as described in Figure 5. **p<0.01

### Figure S5

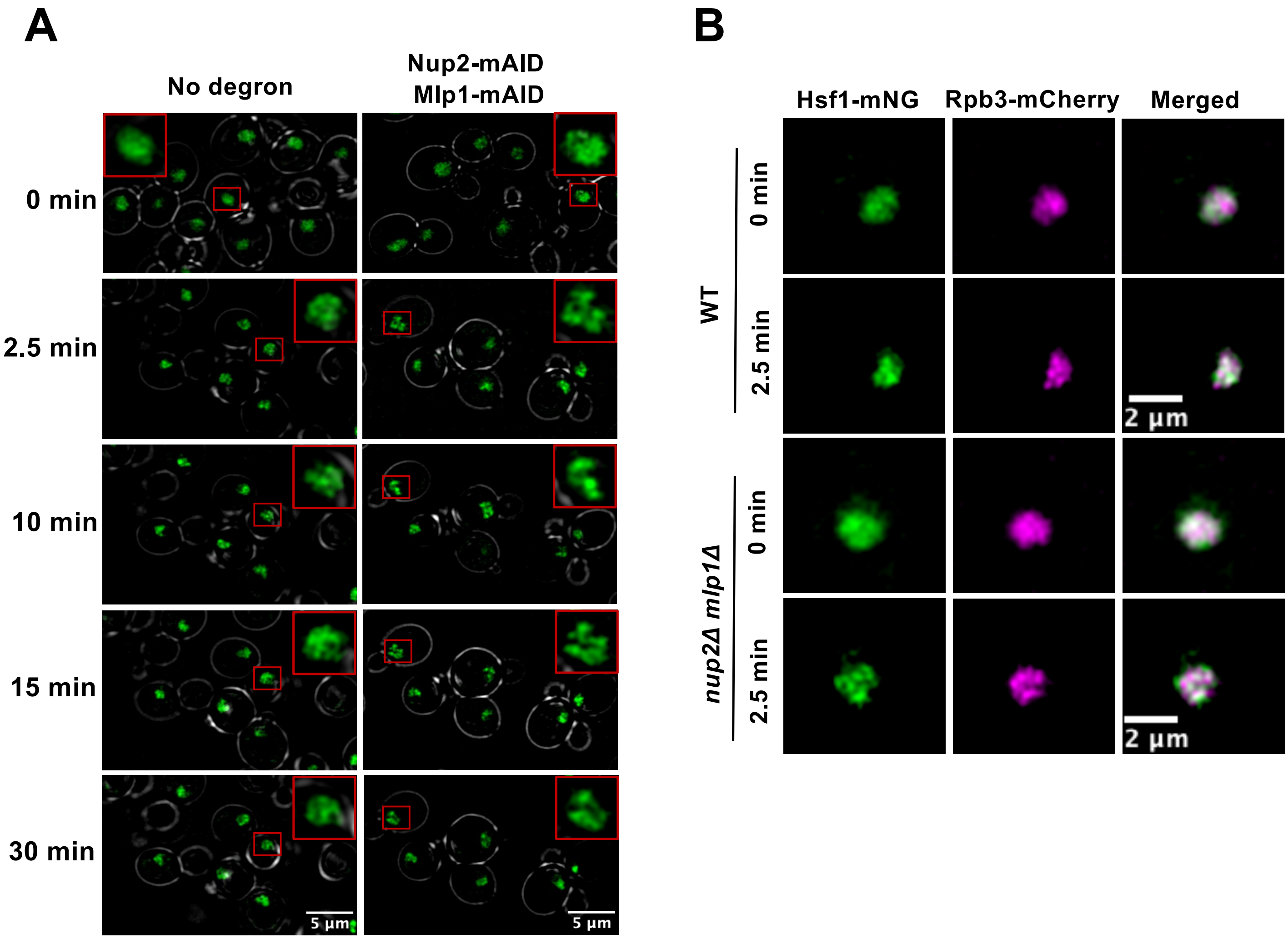


**Figure S5. Formation of transcriptional condensates is unimpeded in Nup2-, Mlp1-depleted cells.**

(A) Hsf1-mNG subnuclear localization in SMY172 (No degron) and SMY170 (Nup2-mAID +Mlp1-mAID) cells treated with auxin for 30 min at 30°C as described in Figure 7A, then subjected to a 39°C HS for the indicated times. A representative fluorescence image of the Z stack is shown for each condition. Z step size: 0.56 microns. Insets are zoomed-in images of the indicated nuclei. See Figure 7B for 3D reconstructions of comparable images.

(B) Subnuclear localization of Hsf1-mNG and Rpb3-mCherry in WT and *nup2Δ mlp1Δ* cells (LRY040 and LRY120, respectively) under NHS (0 min; 25°C) and HS (39°C) conditions as indicated. Depicted are representative Z-planes (step size: 0.5 microns).

Cells in both (A) and (B) were imaged using an Olympus spinning disk confocal microscope. Deconvolution was performed using the Wiener filter plugin in CellSens software.

Figure S6

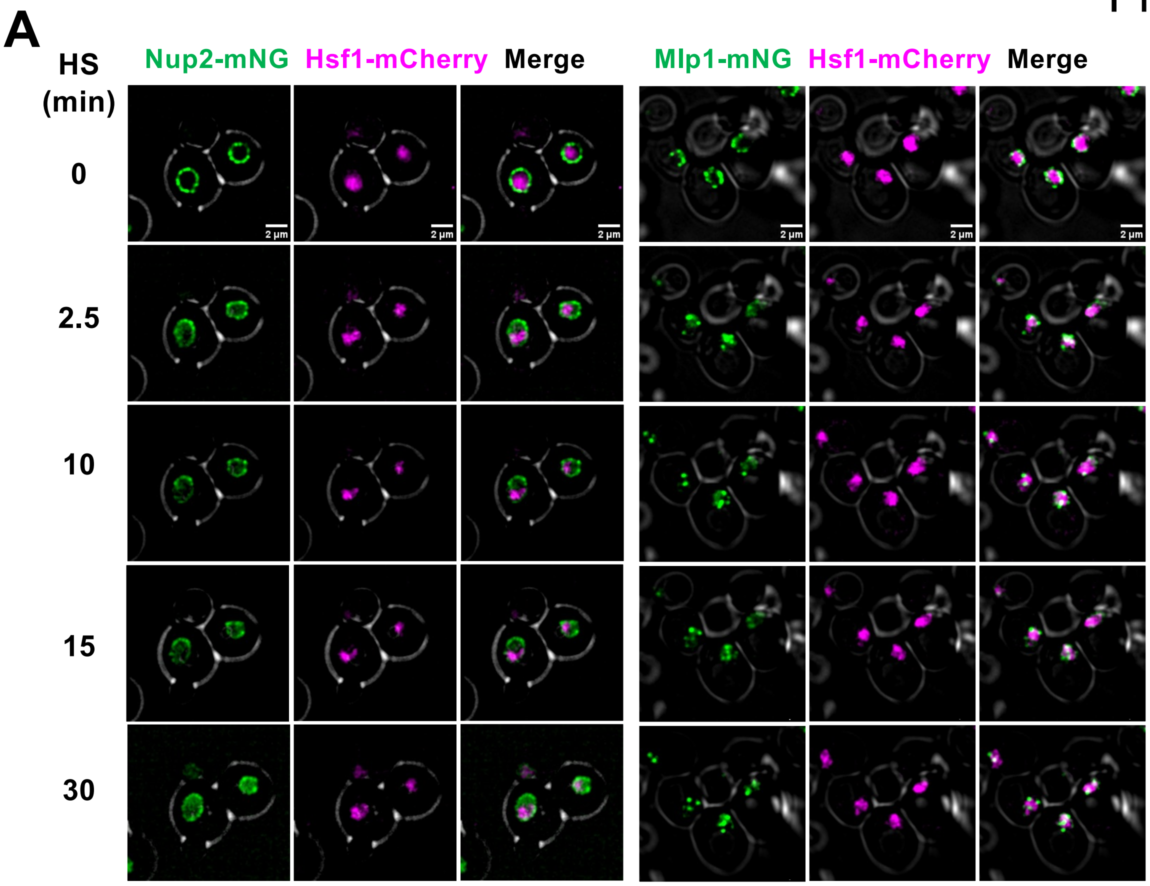


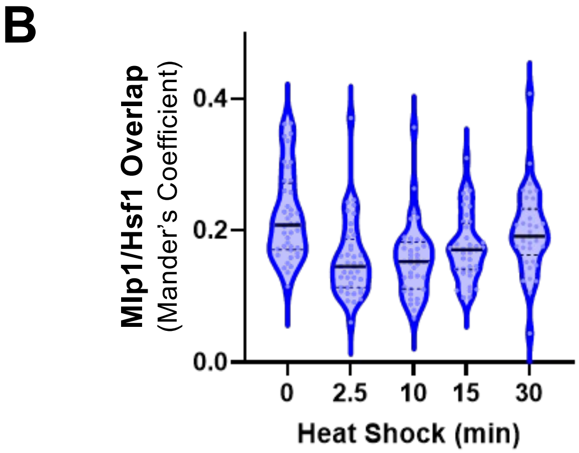


**Figure S6. Nup2 and Mlp1 alter their subcellular distribution upon acute thermal stress yet do not colocalize with Hsf1.**

A. Live cell microscopy of strains LRY888 and LRY777 expressing Hsf1-mCherry in combination with Nup2-mNG and Mlp1-mNG, respectively. Cells were attached to a VAHEAT substrate and imaged before and after thermal upshift (23° to 39°C) for the indicated timepoints. Image acquisition was done using an Olympus spinning disk confocal microscope, with a 0.5 µm interplanar distance over 11 Z-stacks. Images were deconvoluted using Weiner plugin in CellSens software (see Methods). A representative Z plane is shown for each timepoint. Scale bar: 2 µm.

B. Quantification of overlap of Mlp1-mNG and Hsf1-mCherry fluorescence signals within single cells. Strain LRY777 was imaged and subjected to HS as described in (A), followed by determination of Mander's coefficient using the JACoP plugin in ImageJ/FIJI (v. 1.54f) (ref. 116). An average of 70 cells per timepoint derived from two independent cultures are depicted in the violin plots.
