## Supplemental Tables for "Nuclear basket proteins Nup2 and Mlp1 drive heat shock-induced 3D genome restructuring downstream of transcriptional activation"

### Table S1. Yeast Strains

| Strain Name | Genotype | Source |
|---|---|---|
| W303-1A | *MATa ade2-1 trp1-1 can1-100 leu2-3,112 his3-11,15 ura3-1* | R. Rothstein |
| W303-1B | *MATα ade2-1 trp1-1 can1-100 leu2-3,112 his3-11,15 ura3-1* | R. Rothstein |
| ASK701 | *MATα ade2-1 can1-100 leu2-3,112 trp1-1 ura3-1 his3-11,15::GFP-LacI::HIS3 HSP104-LacO*$_{256}$*::TRP1 SEC63-Myc×13::KANMX* | (58) |
| ASK706 | *MATa/MATα ade2-1/ade2-1 can1-100/can1-100 his3-11,15::GFP-LacI::HIS3/his3-11,15::GFP-LacI::HIS3 leu2-3,112/leu2-3,112 trp1-1/trp1-1 ura3-1/ura3-1 HSP12-LacO*$_{128}$*::URA3 /HSP12*$^{+}$ *HSP104-LacO*$_{256}$*:: TRP1 /HSP104*$^{+}$ *SEC63-Myc×13::TRP1 /SEC63-MYC×13::KAN-MX POM34-mCherry::NAT/POM34*$^{+}$ | (17) |
| ASK722 | *MATα ade2-1 can1-100 leu2-3,112 trp1-1 ura3-1 his3-11,15::GFP-LacI::HIS3 HSP104-LacO*$_{256}$*::TRP1 SEC63-MYC×13 TMA10-tetO*$_{200}$*::LEU2* | (17) |
| ASK726 | *MATa ura3-1 thr1-4 ade2-1 leu2-3,112 trp1*$^{-}$ *his*$^{-}$ *leu2::TetR-mCherry::hphMX::leu2* | (17) |
| ASK727 (ASK722 x ASK726) | *MATa/MATα ura3-1/ura3-1 ade2-1/ade2-1 leu2-3,112/leu2-3,112 CAN*$^{+}$ */can1-100 thr1-4 /THR*$^{+}$ *trp1*$^{-}$ */trp1-1 his*$^{-}$ */his3-11,15::GFP-LacI::HIS3 HSP104*$^{+}$ */HSP104-LacO*$_{256}$*::TRP1 TMA10*$^{+}$ */TMA10-TetO*$_{200}$*::LEU2 leu2::TetR-mCherry::hphMX::leu2 SEC63*$^{+}$ */SEC63-Myc×13* | (17) |
| DPY032 | *MAT a ADE2 trp1-1 can1-100 leu2-3,112 his3-11,15 ura3-1 HSF1-mVenus::HIS3* | (18) |
| JTY001 | *MATa ade2-1 can1-100 his3-11,15::GFP-LacI::HIS3 leu2-3,112 trp1-1 ura3-1 HSP12-LacO*$_{128}$*::URA3 HSP104-LacO*$_{256}$*::TRP1 SEC63-MYC×13::KANMX POM34-mCherry::NAT* | (58) |
| LRY016 | *MATα ade2-1 trp1-1 can1-100 leu2-3,112 his3-11,15 ura3-1 LEU2::pGPD1-osTIR1* | (65) |
| LRY037 | W303-1B; *HSF1-mNeonGreen::SpHIS5* | (19) |
| LRY040 | LRY037; *RPB3-mCherry::hphMX6* | (19) |
| LRY116 | LRY040; *mlp1Δ::LoxP-klURA3-LoxP* | This Study |
| LRY120 | LRY116; *nup2Δ::KAN-MX* | This Study |
| LRY117 | SMY125; *mlp1Δ::LoxP-klURA3-LoxP* | This Study |
| LRY118 | SMY127; *mlp1Δ::LoxP-klURA3-LoxP* | This Study |
| LRY119 *(LRY117 X LRY118)* | *MATa/MATα ade2-1/ade2-1 can1-100/can1-100 leu2-3,112/leu2-3,112 trp1-1/ trp1- ura3-1/ura3-1 his3-11,15/his- HSP82*$^{+}$*/HSP82-TetO*$_{<200}$*::LEU2 HSP104*$^{+}$*/HSP104-LacO*$_{256}$*::TRP1 his3::P*$_{REV1}$ *lacI GFP* | This Study |

| | | |
|---|---|---|
| | *$P_{REV1}$ tetR mCherry::HIS3 nup2Δ::KanMX/nup2Δ::KanMX mlp1Δ::LoxP-klURA3-LoxP/mlp1Δ::LoxP-klURA3-LoxP* | |
| LRY777 | SMY192; *HSF1-mCherry::URA3* | This Study |
| LRY888 | SMY216; *HSF1-mCherry::URA3* | This Study |
| SMY108 | ASK726; *HSP82::KANMX* | This Study |
| SMY109 | SMY108; *HSP82-TetO$_{<200}$::LEU2* | This Study |
| SMY110 | *MATa ade2-1 can1-100 leu2-3,112 trp1-1 ura3-1 his3-11,15 HSP104-LacO$_{256}$::TRP1* | This Study |
| SMY118 | *MATα ura3-1 ade2-1 leu2-3,112 trp1$^{-}$ his$^{-}$ HSP82-TetO$_{<200}$::LEU2* | This Study |
| SMY123 | *SMY110; his3::$P_{REV1}$ LacI-GFP $P_{REV1}$TetR-mCherry::HIS3* | This Study |
| SMY125 | SMY123; *nup2Δ::KAN-MX* | This Study |
| SMY127 | SMY118; *nup2Δ::KAN-MX* | This Study |
| SMY134 | DPY032, *nup2Δ:: KAN-MX* | This Study |
| SMY136 | DPY032, *mlp1Δ:: KAN-MX* | This Study |
| SMY143 | LRY016; *NUP1- mAID-Myc9::KAN-MX* | This Study |
| SMY145 | LRY016; *MLP1-mAID-Myc9::KAN-MX* | This Study |
| SMY148 | SMY143; *NUP145-mAID-Myc9::HYG-MX* | This Study |
| SMY149 | LRY016; *NUP2-mAID-Myc9::HYG-MX* | This Study |
| SMY152 | SMY145; *NUP2-mAID-Myc9::HYG-MX* | This Study |
| SMY160 | LRY016; *POM34-mCherry::NAT* | This Study |
| SMY163 | SMY148; *POM34-mCherry::NAT* | This Study |
| SMY164 | LRY016; *NUP2-Myc9::TRP1* | This Study |
| SMY166 | LRY016; *MLP1-Myc9::TRP1* | This Study |
| SMY170 | SMY152; *HSF1-mNeonGreen::HIS5* | This Study |
| SMY172 | LRY016; *HSF1-mNeonGreen::HIS5* | This Study |
| SMY182 | SMY152; *POM34-mCherry::NAT* | This Study |
| SMY192 | LRY016; *MLP1-mNeonGreen::SpHIS5* | This Study |
| SMY196 | SMY148; *MLP1-mNeonGreen:: SpHIS5* | This Study |
| SMY201 | *MATa/MATα ura3-1/ura3-1 ade2-1/ade2-1 leu2-3,112/leu2-3,112 CAN$^{+}$ /can1-100 thr1-4 /THR$^{+}$ trp1$^{-}$ /trp1-1 his$^{-}$ /his3-11,15::GFP-LacI::HIS3 HSP104$^{+}$ /HSP104-LacO$_{256}$::TRP1 TMA10$^{+}$ /TMA10-TetO$_{200}$::LEU2 leu2::TetR-mCherry::hphMX::leu2 SEC63$^{+}$ /SEC63-Myc×13 nup2Δ::KANMX/ nup2Δ::KANMX* | This Study |
| SMY203 | *MATa/MATα ura3-1/ura3-1 ade2-1/ade2-1 leu2-3,112/leu2-3,112 CAN$^{+}$ /can1-100 thr1-4 /THR$^{+}$ trp1$^{-}$ /trp1-1 his$^{-}$ /his3-11,15::GFP-LacI::HIS3 HSP104$^{+}$ /HSP104-LacO$_{256}$::TRP1 TMA10$^{+}$ /TMA10-TetO$_{200}$::LEU2 leu2::TetR-mCherry::hphMX::leu2 SEC63$^{+}$ /SEC63-Myc×13 mlp1Δ::KANMX/ mlp1Δ::KANMX* | This Study |
| SMY206 (SMY118xSMY123) | *MATα/MATa ura3-1/ura3-1 ade2-1/ade2-1 can1-100/ can1-100 leu2-3,112/leu2-3,112 trp1$^{-}$/trp1-1 his$^{-}$ /his3::$P_{REV1}$ LacI-GFP $P_{REV1}$ TetR mCherry::HIS3 HSP82-TetO$_{<200}$::LEU2/HSP82$^{+}$ HSP104$^{+}$/HSP104-LacO$_{256}$::TRP1* | This Study |
| SMY207 | *MATα/MATa ura3-1/ura3-1 ade2-1/ ade2-1 leu2-3,112/leu2-3,112 can1-100 trp1$^{-}$ /trp1-1 his$^{-}$ /his3-11,15 his3::$P_{REV1}$ LacI-GFP $P_{REV1}$ TetR mCherry::HIS3 HSP82-* | This Study |

| | | |
|---|---|---|
| | *$TetO_{<200}$::LEU2/HSP82$^{+}$ HSP104$^{+}$/ HSP104-$LacO_{256}$::TRP1 mlp1Δ::KAN-MX/mlp1Δ::KAN-MX* | |
| SMY208 | *MATα/MATa ura3-1/ura3-1 ade2-1/ade2-1 can1-100 leu2-3,112/ leu2-3,112 trp1$^{-}$/trp1-1 his$^{-}$/his3-11,15 his3::$P_{REV1}$ LacI- GFP $P_{REV1}$ TetR mCherry::HIS3 HSP82-$TetO_{<200}$::LEU2/HSP82$^{+}$ HSP104$^{+}$ /HSP104-$LacO_{256}$::TRP1 nup2Δ::KAN-MX/nup2Δ::KAN-MX* | This Study |
| SMY216 | LRY016; *NUP2-mNeonGreen:: SpHIS5* | This Study |
| SMY221 | SMY148; *NUP2-mNeonGreen:: SpHIS5* | This Study |

### Table S2. Plasmids

| Plasmid Name | Feature | Source |
|---|---|---|
| pSR14 | LEU2-$TetO_{200}$ array | Susan Gasser (117) |
| pFA6a-kanMX6 | pFA6a-kanMX6 | Addgene Plasmid #39296 (118) |
| pFA6a-link-ymNeonGreen-SpHis5 | link-mNeonGreen-SpHIS5 | (119) |
| pWZV87 | Myc9-KITRP1 | Kim Nasmyth (120) |
| pMY63 | REV1pr-LacI-GFP-REV1pr-TetR-mCherry, HIS3 marker | Lu Bai (121) |
| pHyg-AID*-9myc | Mini AID-9MYC-HYGR | Addgene Plasmid #99518 (98) |
| pKAN-AID*-9myc | Mini AID-9MYC-KANMX | Addgene Plasmid #99522 (98) |
| pUG72 | LoxP-KIURA3-LoxP | (122) |

### Table S3. Primers used for Strain Construction

| Name | Sequence (5' → 3') | Purpose |
|---|---|---|
| *HSP82_KANMX_* Chimeric_F | GTTATAAACAAAACATAATATAACGTATAGGTATTC GAATGAATAAATAAAGCGGATGCCGGGAGCAGAC | Insertion of *KAN-MX* into the 3' end of *HSP82* |
| *HSP82_KANMX_* Chimeric_R | ATTGTAATGTTTTACCCAGTTATTTCCATGCAGATG CCCTATTTACATACGTGAGCTGATACCGCTCGCC | |
| *HSP82*_Conf 1_F | AGCTGACACCGAAATGGAAGAGG | Confirmation of *KAN-MX* insertion |
| *HSP82*_Conf 1_R | GTTGGACGCATAATGAAAGCAGATGAG | |
| *HSP82* _Conf 2_F | AGCTGACACCGAAATGGAAGAGG | To confirm the insertion of the *LEU2-TetO* array at the *KAN-MX* locus |
| *HSP82*_Conf 2_R | AGCAGACAAGATAGTGGCGATAGGG | |
| *NUP145*-C-HYG-AID-F | TGAGTTTGCCCAGGATTTAATGAAGTGTACATATA AGATACGTACGCTGCAGGTCGAC | *NUP145* - degron C-term tagging |
| *NUP145*-C-HYG-AID-R | CCATGTTTTACTATTTTTCTTTTTTTTAGAAATAAAA ATAAAAAAACTCGATGAATTCGAGCTCG | |
| *NUP145*_C-term_conf_ F | GATCAGTATAAGCACTGTCGTGAAGTGG | Confirmation of *NUP145*-C-term tagging |
| *NUP145*_C-term conf_R | GAGGATTGGCAAGAGTTGTGACATGGG C | |
| *NUP1*-C-KAN-AID-F | TGGCGAACAGAAAGATTGCAAGAATGAGGCACTC TAAAAGGCGTACGCTGCAGGTCGAC | *NUP1*-degron C-term tagging |
| *NUP1*-C-KAN-AID-R | CCTTCAGAAAAGCAACACAATACCTAATTACATAA CCGATATTCGATGAATTCGAGCTCG | |
| *NUP1*_C-term conf_F | CAGTCGTCACTCATGGTGATTTCTCAC | Confirmation of *NUP1*-C-term tagging |
| *NUP1*_C-term conf_R | CCATCATTGTTGTGAATACGCACCC | |
| *NUP2*-C-HYG-AID-F | CTCATTTACGAAAGCTATTGAAGATGCTAAAAAAG AAATGAAACGTACGCTGCAGGTCGAC | *NUP2*-degron C-term tagging |
| *NUP2*-C-HYG-AID-R | AGGGTTCTATTCTATTTAAAATTGTTAACTGTATTT ACTCTCGATGAATTCGAGCTCG | |
| *NUP2*_C-term conf_F | ATGTGTATCACTGGCAAACTGTGATGG | Confirmation of *NUP2*-C-term tagging |
| *NUP2*_ C-term conf_R | AGGGATGAAAGAAGATTGGCTTGGG | |
| *MLP1*-C-KAN-AID-F | GGAAGAAAAAGAAACCGATAAGGTGAATGACGAG AACAGTATACGTACGCTGCAGGTCGAC | *MLP1*-degron C-term tagging |
| *MLP1*-C-KAN-AID-R | AGGTTTAGTTTGTATTGATCCCTTGTTTTTACTATC TCCTTCGATGAATTCGAGCTCG | |
| *MLP1*_C-term conf_F | CAGTCGTCACTCATGGTGATTTCTCAC | Confirmation of *MLP1*-C term tagging |
| *MLP1*_C-term conf_R | TGACATAGGGCAGAATGAAGCTCCTCC | |
| *POM34*-mCherry-F | GCTCTTATCCACCGTCAAAGTAAGTG | |

| | | |
|---|---|---|
| *POM34*-mCherry-R | CAAATCCTGAATCCGAAGAACCGTGC | Tagging *POM34* with mCherry |
| *POM34*-mCherry_conf_F | CACACCACGTTCAGTTGGTTGAATGC | Confirmation of *POM34*-mCherry tagging |
| *POM34*-mCherry_conf_R | TCCTGTCACAATCTCTCAGTTCGTAGG | |
| *HSF1*-ymNeonGreen F | CGAGAACGCTAAGAAAAGATTTGTGG | Tagging *HSF1* with mNeonGreen |
| *HSF1*-ymNeonGreen R | GTGCAGTTCAACCTCACTCG | |
| *HSF1*_conf_F | TGACCACAGTTATTCCACC | Confirmation of *HSF1*-mNeonGreen tagging |
| *HSF1*_conf_R | CCAATGTGACACCAGTTCACTCG | |
| *NUP2*-C-9Myc_F | CTCATTTACGAAAGCTATTGAAGATGCTAAAAAAG AAATGAAATCCGGTTCTGCTGCTAG | Tagging *NUP2* with 9Myc |
| *NUP2*-C-9Myc_R | AGGGTTCTATTCTATTTAAAATTGTTAACTGTATTT ACTCCCTCGAGGCCAGAAGAC | |
| *NUP2*_Conf_ F | GGAAGAATCAACAACAGAAGCAACTGG | Confirmation of *NUP2*-C-term 9Myc tagging |
| *NUP2*_Conf_ R | AGGGATGAAAGAAGATTGGCTTGGG | |
| *MLP1*-C-9Myc_F | GGAAGAAAAAGAAACCGATAAGGTGAATGACGAG AACAGTATATCCGGTTCTGCTGCTAG | Tagging *MLP1* with 9Myc |
| *MLP1*-C-9Myc_ F | AGGTTTAGTTTGTATTGATCCCTTGTTTTTACTATC TCCTCCTCGAGGCCAGAAGAC | |
| *MLP1*_conf_ F | CATCGAACAGAAATGTTCAATCGGAAGAG | Confirmation of *MLP1*-C term-9Myc tagging |
| *MLP1*_conf_ R | TGACATAGGGCAGAATGAAGCTCCTCC | |
| *NUP2*_F | GTGGTAAACAAGCTTCCACCGAATGAG | To replace *NUP2* ORF with *KAN-MX* |
| *NUP2*_R | GATAGGGATGAAAGAAGATTGGCTTGGG | |
| *NUP2*_conf_F | ACAGTAGCACATCCGTGAAACTTCTGG | Confirmation of *NUP2* deletion |
| *NUP2*_Kan specific conf_R | GAGTAACCATGCATCATCAGGAGTACGG | |
| *MLP1*_F | TGACTAGGACTTAACTGATACTCGCCGAAG | To replace *MLP1* ORF with *KAN-MX* |
| *MLP1*_R | TAGGGCAGAATGAAGCTCCTCCACATTG | |
| *MLP1*_conf _F | GGGATAGATGGGTAATGGCTAGTATGAGGC | Confirmation of *MLP1* deletion |
| *MLP1*_kan specific conf_R | GAGTGACGACTGAATCCGGTGAGAATGG | |
| *MLP1*-C-ymNeonGreen_F | GGAAGAAAAAGAAACCGATAAGGTGAATGACGA GAACAGTATA GGTGACGGTGCTGGT | Tagging *MLP1* with mNeonGreen |
| *MLP1*-C ymNeonGreen_R | AGGTTTAGTTTGTATTGATCCCTTGTTTTTACTAT CTCCTTCGATGAATTCGAGCTCG | |
| *MLP1* conf _F | CATCGAACAGAAATGTTCAATCGGAAGAG | Confirmation of *MLP1*-C term-mNG tagging |
| *MLP1*_conf_R | TGACATAGGGCAGAATGAAGCTCCTCC | |

| | | |
|---|---|---|
| *MLP1 del F* | GATACTCGCCGAAGCTACACAAATAGTCAGTAACGCCACGTTTTAGGATACAGCTGAAGCTTCGTACGC | To delete *MLP1* with LoxP system |
| *MLP1 del R* | ACATTGAAAAAGGTTTAGTTTGTATTGATCCCTTGTTTTTACTATCTCCTGCATAGGCCACTAGTGGATCTG | |
| *MLP1* del Conf_F | GCAAATTGAATACAGACAGAGATATC | Confirmation of *MLP1* deletion |
| *MLP1* del Conf_R | CTATTTACGTGACTTCATCTTAGC | |
| *NUP2*-C-ymNeonGreen_F | CTCATTTACGAAAGCTATTGAAGATGCTAAAAAAGAAATGAAAGGTGACGGTGCTGGT | Tagging *NUP2* with mNeonGreen |
| *NUP2*-C-ymNeonGreen_R | AGGGTTCTATTCTATTTAAAATTGTTAACTGTATTTACTCTCGATGAATTCGAGCTCG | |
| *NUP2*_conf _F | GGAAGAATCAACAACAGAAGCAACTGG | Confirmation of *NUP2*-C term-mNG tagging |
| *NUP2*_conf_R | AGGGATGAAAGAAGATTGGCTTGGG | |
| *HSF1*-C-term mCherry_F | AGGACCCGACAGAGTACAACGATCACCGCCTGCCCAAACGAGCTAAGAAAGGTGACGGTGCTGGTTTA | Tagging of Hsf1 with mCherry |
| *HSF1*-C-term mCherry_R | ATACTATATTAAATGATTATATACGCTATTTAATGACCTTGCCCTGTGTATCGATGAATTCGAGCTCG | |
| *HSF1*_Conf _F | ACGACAATAACACTAGTGAGG | Confirmation of Hsf1-mCherry tagging |
| *HSF1*_Conf_R | CTCAGGCTCTCACTAGCTC | |

### Table S4. Primers used for RT-qPCR

| Name | Sequence (5' → 3') |
|---|---|
| *HSP104 ORF F+1646* | CAGCTGCAAGATTGACTGGTATCC |
| *HSP104 ORF R+1799* | CCTGATCTAGACAATCTAACGGC |
| *HSP82 ORF F+290* | CAAGTCTGGTACCAAAGC |
| *HSP82 ORF R+453* | CAGTGAAAGAACCACCAGC |
| *SSA4 ORF F+815* | GTCTTCGTCTGCTCAGACATC |
| *SSA4 ORF R+946* | CCACTGGCTCCAATGTAGATC |
| *HSP12 ORF F+183* | AAAAGGCAAGGATAACGCTGAAG |
| *HSP12 ORF F+327* | CTTCTTGGTTGGGTCTTCTTC |
| *BTN2 ORF F+555* | GTTTTTGTTATTGGCTGTGGAG |
| *BTN2 ORF R+649* | CTTCCTCATGCTTAACTAAACC |
| *SCR1 F+385* | CGGCCGGGATAGCACATATC |
| *SCR1 R+438* | CGCCGAAGCGATCAACTTG |

### Table S5. Primers used for ChIP

| **Name** | **Sequence (5' → 3')** |
|---|---|
| *ARS504 F* | GTCAGACCTGTTCCTTTAAGAGG |
| *ARS504 R* | CATACCCTCGGGTCAAACAC |
| *HSP104 UAS F -266* | CTTAAACGTTCCATAAGGGGC |
| *HSP104 UAS R -195* | TGCAGTTCTTTGAGATGGGCC |
| *HSP104 Prom F -130* | GCATTGTAATCTTGCCTCAATTCC |
| *HSP104 Prom R -70* | GTTATTGCTGATTCGATTCAAGG |
| *HSP104 ORF F +1469* | CCCTTGATGCTGAACGTAGATATG |
| *HSP104 ORF R +1621* | CCACATTTTGGATCATGGAGTTG |
| *HSP104 3'UTR F +2676* | AGGTGATGACGATAATGAGGACAG |
| *HSP104 3'UTR R +2839* | TCTTTTGCTCGGGTGTCAAGTTC |
| *HSP82 Prom F -157* | TCCGCCACCCCCTAAAAC |
| *HSP82 Prom R -113* | TGAGGAGGTCACAGATGTTAAGAATT |
| *HSP82 ORF F +1392* | GCCAGAACACCAAAAGAACATCTAC |
| *HSP82 ORF R+1522* | ATTCATCAATTGGGTCGGTCAAG |
| *HSP82 3'UTR F +2036* | ATGAGGATGAAGAAACAGAGACTGC |
| *HSP82 3'UTR R+2297* | ACACACTAGACGCGTCGGAATAG |
| *SSA4 UAS F -374* | GCCGCACATCCATTCCGGTATG |
| *SSA4 UAS R -291* | CGGGCAAAAGATATCCGCTTTG |
| *SSA4 Prom F -246* | AGTTCCTAGAACCTTATGGAAGCAC |
| *SSA4 Prom R +35* | GTTGTACCTAAATCAATACCAACAGC |
| *SSA4 ORF F +816* | GTCTTCGTCTGCTCAGACATC |
| *SSA4 ORF R +946* | CCACTGGCTCCAATGTAGATC |
| *SSA4 3'UTR F +1762* | GAGGAATACAAGGAAAGGCAAAAG |
| *SSA4 3'UTR R +2079* | TTAAACTCTGGCTTATGACGATGAG |

### Table S6. Primers used for Taq I-3C

| Name | Sequence (5' → 3') |
|---|---|
| *ARS504 F* | GTCAGACCTGTTCCTTTAAGAGG |
| *ARS504 R* | CATACCCTCGGGTCAAACAC |
| *HSP12 F-47* | ACGTATAAATAGGACGGTGAATTGC |
| *HSP12 R-47* | TTCAGAAGCTTTTTCACCGAATC |
| *HSP82 F+740* | AATTAGTCGTCACCAAGGAAGTTG |
| *HSP82 R+740* | AATGCTTAACGTACAATGGGTCTTC |
| *HSP82 F+2189* | ATGAGGATGAAGAAACAGAGACTGC |
| *HSP82 R+2189* | ACACACTAGACGCGTCGGAATAG |
| *HSP104 F-63* | AGGCATTGTAATCTTGCCTCAATTC |
| *HSP104 R-63* | ATCGTTAGAGCCCTTTCTGTAAATTG |
| *HSP104 F+782* | GTAAGACCGCTATTATTGAAGGTG |
| *HSP104 R+782* | TTCTTCGATTTCCTTCAAAACACC |
| *HSP104 F+1550* | CCCTTGATGCTGAACGTAGATATG |
| *HSP104 R+1550* | CCACATTTTGGATCATGGAGTTG |
| *HSP104 F+2756* | AGGTGATGACGATAATGAGGACAG |
| *HSP104 R+2756* | TCTTTTGCTCGGGTGTCAAGTTC |
| *SSA2 F+198* | AGGTAACAGAACCACTCCATCTTTC |
| *SSA2 R+198* | GCTTCATATCACCTTGGACTTCTG |
| *SSA2 F+1368* | TCTCTACTTATGCTGACAACCAACC |
| *SSA2 R+1368* | TTCAATTTGTGGGACACCTCTTG |
| *SSA4 F-268* | ACACGAAAGATATCTCAACTCTAGCC |
| *SSA4 R-268* | TGTTACTGTCGTCAAACTAAGGAG |
| *SSA4 F+198* | GCCTTCTTATGTGGCTTTTACTGAC |
| *SSA4 R+198* | TTTACGTCCGATCAGACGCTTAG |
| *SSA4 F+1079* | TGCTGATTTGTTTAGATCTACATTGG |
| *SSA4 R+1079* | TAATACCACCTGCAGTTTCAATACC |
| *SSA4 F+2255* | ATAAGAAAGTCATCGCCAAACAAC |
| *SSA4 R+2255* | GTGTTAAACTCCGGTCAAAAGAAAC |
| *UBI4 F+524* | GTAAGCAGCTAGAAGATGGTAGAACC |
| *UBI4 R+524* | TGAATTTTCGACTTAACGTTGTCG |
